# Efficient self-organization of blastoids solely from mouse ESCs is facilitated by transient reactivation of 2C gene network

**DOI:** 10.1101/2023.04.12.536583

**Authors:** Debabrata Jana, Priya Singh, Purnima Sailasree, Nithyapriya Kumar, Vijay V Vishnu, Hanuman T Kale, Jyothi Lakshmi, Asha Kumari, Divya Tej Sowpati, P Chandra Shekar

## Abstract

Human pluripotent stem cells (hPSCs) can self-organize into a blastocyst-like structure (blastoid) by virtue of their full developmental potential. The pluripotent mouse embryonic stem cells (mESC) are considered to lack this potential and hence can form blastoids only when combined with trophoblast stem cells. We performed a small molecule and cytokine screen to demonstrate that mESC have full potential to efficiently self-organize themselves into E-blastoids (ESC-blastoids). The morphology, cell lineages and the transcriptome of these blastoids resemble the mouse blastocyst. The E-blastoids undergo implantation and *in utero* development in mice. The transient reactivation of the 2C-gene network by retinoid signaling is essential for E-blastoid generation. GSK3β activity is critical for retinoid signaling and consequent 2C gene network activation. Collectively, the mESC possess full developmental potential to generate blastoids similar to hPSCs and other mammals. The plasticity of PSCs to self-organize into blastoids is not exclusive to humans or larger mammals; rather, it could be a general feature shared by most mammals, including rodents.

## Introduction

Murine embryonic stem cells (ESCs) cultured in Serum and LIF (SL) are pluripotent and have characteristic features of the epiblast of late blastocyst(Nichols et al., 2009). They exist in a state of metastability, transiently giving rise to subpopulation cells with robust developmental potential similar to the 2-cell stage blastomeres and ground state pluripotent stem cells at low frequency(Macfarlan et al., 2012; Ying et al., 2008). Multiple stem cells with robust developmental potential have been derived from ESCs or cleavage stage embryos, including expanded or extended pluripotent stem cells (EPSCs), totipotent blastomere-like cells (TBLCs), totipotent-like stem cells (TLSCs), and totipotent potential stem cells (TPS) with trophoectoderm potential(Shen et al., 2021; Xu et al., 2022; Yang et al., 2017a; Yang et al., 2022; Yang et al., 2017b; Zhang et al., 2022). These cells have the potential to contribute to all three layers of the blastocyst, unlike the ESCs.

Stem cell lines derived from mammalian embryos can organize *in vitro* into blastocyst-like structures called blastoids, which structurally resemble natural embryos and can implant and undergo post-implantation development until early to mid-gestation. Blastoids are composed of all cell types of the blastocyst: trophoectoderm, epiblast, and primitive endoderm. Murine blastoids were first generated by combining ground state (2iL) ESCs with trophoblast stem cells (TSCs), where the ESCs contributed to the epiblast and the primitive endoderm, while the TSCs contributed to the trophoectoderm layer of the blastoids(Rivron et al., 2018). Blastoids have also been generated from a single type of stem cell with robust developmental potential in absence of TSCs(Li et al., 2019; Shen et al., 2021; Xu et al., 2022; Zhang et al., 2022).

hPSCs have unrestricted lineage potential and readily differentiate to trophoectoderm(Guo et al., 2021). The full developmental plasticity of hPSCs has enabled the generation of blastoids exclusively from hPSC at high efficiency in absence of TSCs(Shen et al., 2021; Xu et al., 2022; Zhang et al., 2022). Recently, blastoids have also been generated solely from naive ESCs of monkeys(Li et al., 2023). However, blastoids can only be produced when TSCs are coupled with mouse ESCs; they cannot be generated from mESCs alone. When ESCs alone have been used to produce blastoids, a *Cdx2* transgene was activated in the ESC to produce the trophoectoderm compartment of blastoids(Lau et al., 2022). The lack of trophoectoderm potential is considered to limit the ability of the ESC to self-organize into the blastoids by themselves. The unrestricted potential of hPSCs is attributed to the lack of chromatin-based lineage barriers in hPSCs. In contrast, it was recently reported that the histone posttranslational modifications (hPTMs) in naive hPSCs and mouse ESCs are largely conserved and hPSC also have chromatin-based lineage barriers(Zijlmans et al., 2022). Surprisingly, blastoids have not been generated solely from mESCs, despite the fact that hPTMS are mostly preserved between naive hPSCs and mESCs.

We asked, whether the mESC possess full potential to self-organize themselves into blastoids. To investigate this, a small-scale screen of small molecules and cytokines was carried out, which resulted in the generation of blastoids from mESCs alone under defined culture conditions without any transgene expression. The blastoids derived from mESCs undergo implantation, induce decidualization, and develop live post-implantation tissues *in utero*. We further show that this inherent full potential of the mESC could be unlocked by transient activation of the 2C gene network during self-organization of mESC to blastoids. GSk3β activity and retinoid signaling were found to be essential for this process. We show that despite their similarity to epiblast of late blastocyst, the mESC possess full potential to self-assemble into blastoids comprising all three embryonic and extraembryonic lineages of blastocysts.

### ESCs efficiently self-organize into blastocyst-like structures

We performed a screen of combinations of cytokines and small molecules to generate blastoids from ESCs (Figure S1A). The ESCs were either cultured in SLPD or SLCHIR or SL for at least 48hrs. Cells were cultured in suspension in various combinations of cytokines and small molecules to aggregate and organize into blastocyst-like structures. These cultures were analyzed at every 24hrs intervals for blastocyst-like structures. They were scored for two parameters – spherical shape and formation of a cavity containing inner cells. We used the culture conditions used for the generation of B-blastoid from EPSCs as a control(Li et al., 2019). Blastocyst-like structures with cavities were observed by 72hrs in some of the culture conditions, where the ESCs were pre-cultured in SLPD for 48hrs. Intriguingly, in a cocktail containing Retinoic Acid (RA) and BMP4 (RB), 61% of the spheres with a cavity resembling blastocysts were observed. This improved to 68% with the addition of bFGF (RBF) (Figure S1A). The efficiency of blastoid organization from ESC using RBF was higher than the previously reported methods using stem cells like EPSCs (12%), and TLSCs (35%)(Li et al., 2019; Sozen et al., 2021). To generate uniform and consistent blastoids, we generated them in Aggrewells with a specific initial number of cells in RBF. Aggregates beginning with around 40 ESCs developed into uniform-sized blastoid structures comparable to blastocysts (Figure S1B). In most of the spheres, a small cavity appeared around 48hrs of self-organization. We supplemented RBF with CHIR at 48hrs to improve the cavity formation (Figure S1C). Together, we developed an efficient method (>90%) to self-organize blastoids exclusively from ESCs (Figure 1A). The blastoids derived from ESCs were referred to as E-blastoids (Ebl) and the method as E-blastoid method (Figure 1A). We analyzed the Ebls for the three embryonic and extraembryonic blastocyst lineages by immunostaining. The outer cells surrounding the cavity expressed CDX2. The ICM-like compartment comprised of cells that expressed EPI marker OCT4 and PE marker GATA4 (Figure 1B). The CDX2, OCT4 and GATA4 expressing cells were localized to the respective blastocyst compartments in E-blastoids. We also generated blastoids from EPSCs as described by Li et al, which will be referred to as B-blastoids (Belmonte lab-blastoids). We analyzed the proportion of the cells in embryonic and extraembryonic lineages in E- and B-blastoids. The number of CDX2 expressing TE lineage cells in E-blastoids were comparable to E3.5 blastocyst, unlike the B-blastoids which had fewer cells (Figure 1C). The E-blastoids had fewer EPI lineage cells. The GATA4 expressing PE lineage cells were 2-3-fold higher in both E- and B-blastoids (Figure 1C). Although all three lineages of blastocyst were detected in Ebl, the EPI compartment was slightly compromised and the PE compartment was expanded. The B-blastoids showed a compromised TE compartment and expansion of EPI and PE compartments (Figure 1B and 1C) suggesting both blastoids show a stochiometric deviation of the three lineages from the blastocyst. These results suggest that the mESC which are *in vitro* counterparts of E4.5 epiblast possess full potential to self-organize into blastocyst like structures containing both embryonic and extraembryonic lineages.

**Figure. 1:**
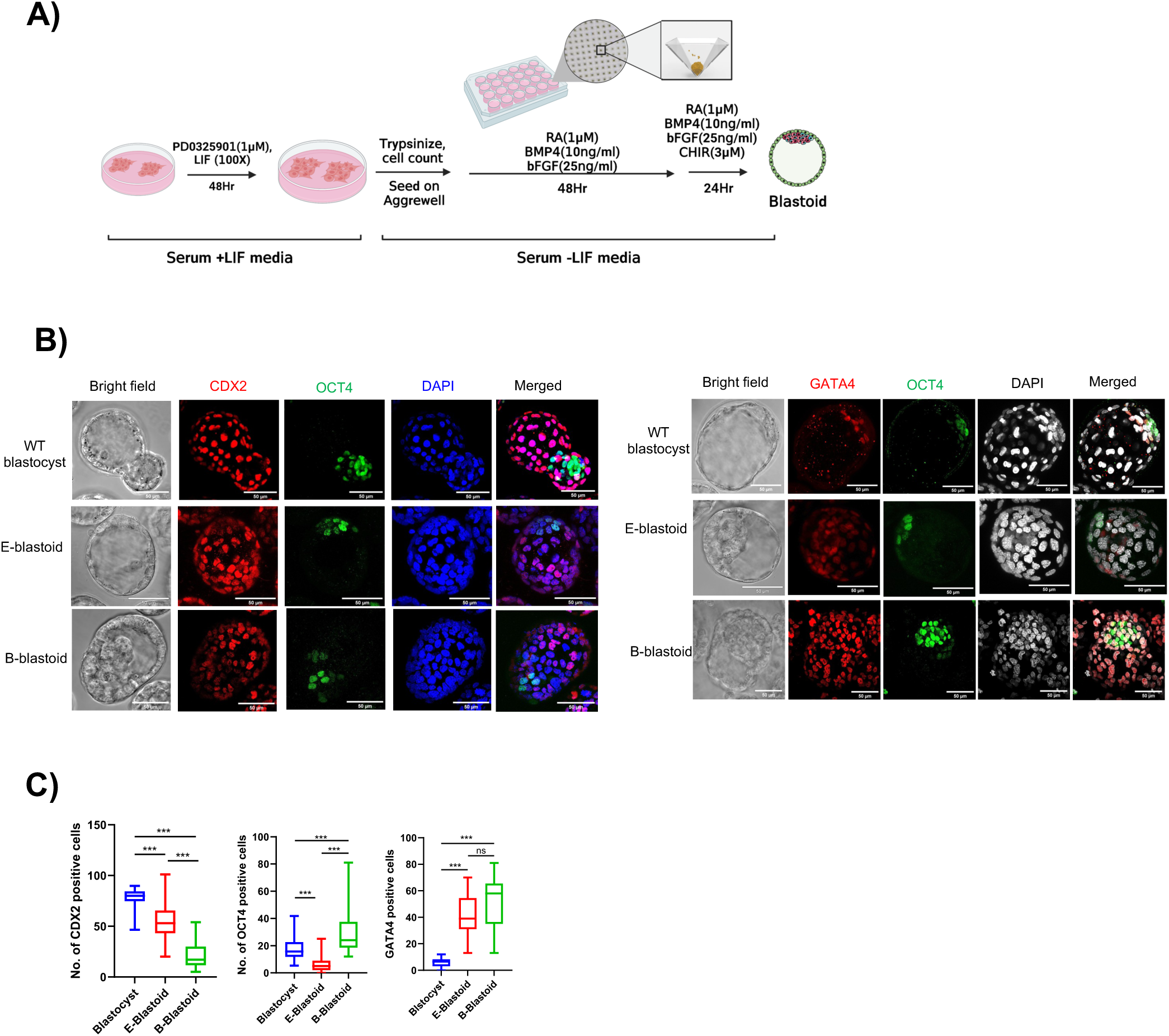
Generation and charcterisation of blastoids from ESCs. (A) Schematic depicting the method followed to generate E-blastoids from ESCs. B) (left) Immunostaining of CDX2, OCT4 in blastocyst, E-blastoid, and B-blastoid. (Right) Immunostaining of GATA4, OCT4 in blastocyst, E-blastoid, and B-blastoid. (C) Quantification of the number of CDX2, OCT4, and GATA4 expressing cells in the blastocyst, E-blastoid, and B-blastoid (n>30). ‘*’ indicates a p-value <0.05, ‘**’ indicates a p-value <0.01, ‘***’ indicates a p-value <0.001, ‘ns’ indicates a p-value > 0.05.

### Functional Characterization of E-blastoids

The ESC, trophoblast stem cells (TSC) and extraembryonic endoderm (XEN) cells are the *in vitro* equivalents of epiblast, trophoectoderm and primitive endoderm of blastocyst respectively(Niakan et al., 2013; Tanaka, 2006; Ying et al., 2008). We could establish and maintain ES-, XEN- and TS-like cell lines from the E-blastoids (Figure 2A, 2B). The ES-like cells were derived in ground state culture condition (2iL). The ES-like cells maintained in 2iL and SL media for multiple passages morphologically resembled ESCs (Figure 2B). They expressed the pluripotency markers and could differentiate to all three germ layers invitro (Figure S2A). The ES-like cells contributed to chimerism and participated in embryonic development, when injected into the blastocyst (Figure S2B). The XEN-like cells morphologically resembled the XEN cells from blastocysts and expressed the XEN marker GATA4 (Figure 2B, S2C). We could derive TS-like cells from E-blastoids in FAXY media (Ohinata and Tsukiyama, 2014) but not in TSC media(Tanaka, 2006) or human TSC media(Okae et al., 2018). The TS-like cells derived from E-blastoids could continuously self-renew and expressed the TSC markers like CDX2 and GATA3 (Figure S2D).

**Figure. 2:**
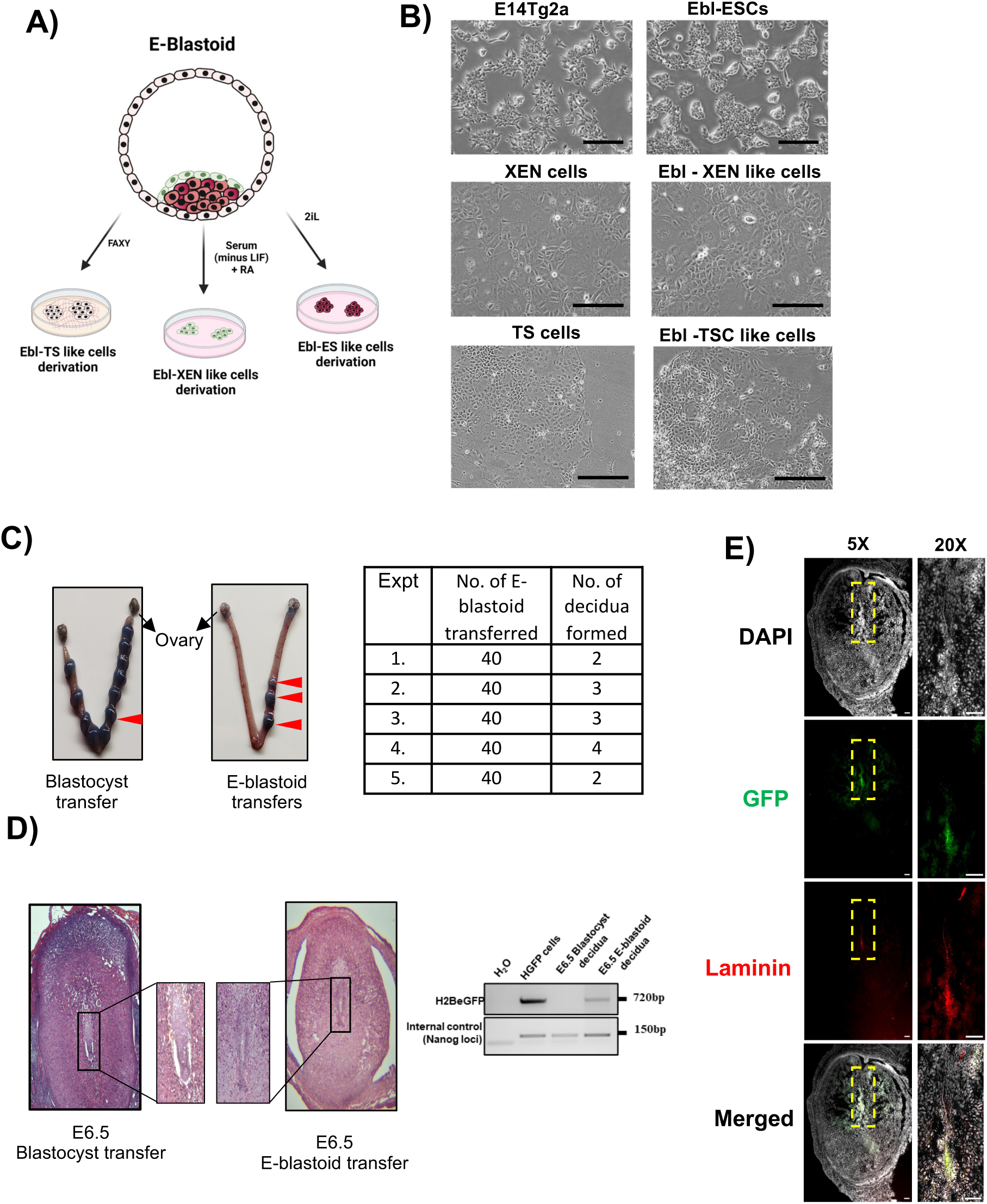
Functional Characterization of E-blastoids. (A) Schematic representation depicting derivation of TS-like cells, XEN-like cells and ES-like cells from E-blastoid. (B)(Top) Phase contrast images of E14Tg2a and ESCs derived from Ebl cultured in SL. (Middle) Phase contrast images of XEN cells (derived from blastocyst) and XEN-like cells derived from Ebl. (Bottom) Phase contrast images of TS cells (derived from blastocyst) and TSC-like cells derived from Ebl. Scale bar = 100µm. (C) (left) Pontamine sky blue staining of the implantation points of blastocyst and blastoid transfers uterus at E6.5. Red arrows indicate the decidualization points. (Right) table summarizing the number of blastoid transfers and the number of deciduae formed. (D) (left) Hematoxylin/Eosin staining of E6.5 deciduae section developed from blastocyst and E-blastoid. (right) PCR amplification of H2B-GFP cassette from the decidua of blastocyst and E-blastoid generated from HGFP cells. (E) Immunostaining for GFP and Laminin in E6.5 decidual sections of E-blastoid transfer. Scale bar = 1000µm.

To further analyse *in vivo* developmental potential of the Ebls to implant and undergo foetal development, we transferred Ebls into the uterus of 2.5 days post coitus (dpc) pseudo-pregnant mice. The post-implantation development was analyzed at E6.5. ESC line with constitutive eGFP expression was utilised in these experiments. The Ebls formed decidua similar to the blastocysts as analyzed by pontamine sky blue staining (Figure 2C). Around 7% implantation efficiency of Ebls was comparable to the implantation efficiencies reported for ETS-blastoids and B-blastoids(Li et al., 2019; Posfai et al., 2021). The establishment of the embryonic axis was evident in the blastocyst and Ebl decidua sections at E6.5 (Figure 2D). The vascular sinuses containing RBCs further confirmed the maternal blood supply (Figure 2D). The PCR of amplification of GFP transgene from genomic DNA of the decidua reconfirmed the contribution of the E-blastoids to the decidua (Figure 2D). Further immunostaining of the deicidal sections revealed presence of the GFP expressing cells co-stained with Laminin suggesting development of the blastoids to the post implantation structures. However, the post-implantation embryonic structures derived from E-blastoids were malformed to varying degrees suggesting developmental defects (Figure 2E). Together these results imply that the E-blastoids can implant, induce decidualization, and progress through *in utero* development since they are made up of cells from all three embryonic and extraembryonic layers of the blastocyst.

### Single-cell Transcriptome analysis of E-blastoids

The E-blastoids were self-organized in approximately 72 hours, which corresponds to the mouse preimplantation development timelines after the first cleavage. We performed single-cell RNA Seq (ScRNA Seq) of E-blastoids at 24hrs (Ebl24), 48hrs (Ebl48) and 72hrs (Ebl) during Ebl generation and compared the transcriptome to the different developmental stages and other b(Posfai et al., 2021)lastoids. The three stages of E-blastoids clustered with the B-blastoids and ZG-blastoids(Sozen et al., 2019) (Zernica-Goetz lab blastoids) in principle component analysis (PCA), suggesting the E-blastoids resembled the blastoids generated using other methods and stem cells (Figure 3A). Notably, all the blastoids clustered distinctly from pre-implantation developmental stages suggesting they are significantly different from pre-implantation stages. The formative ESC (FTW-ESC) was relatively closer to all the blastoids suggesting that blastoids may relate to embryonic stages after E3.5 (Figure 3A). We further analyzed the correlation between each of the blastoids and the developmental stages (Figure 3B). All the stages of E-blastoids showed significant correlation with ZG-blastoids and B-blastoids. All three blastoids showed a good correlation with E5.5 EPI and the PE suggesting, all blastoids irrespective of the starting cell types or methods used for their generation may relate to a developmental stage beyond E3.5. The TE lineage appears to be compromised in all blastoids (Figure 5B). None of the blastoids show any correlation with ICM, further confirming that the blastoids do not represent developmental stages earlier than E4.5.

**Figure. 3:**
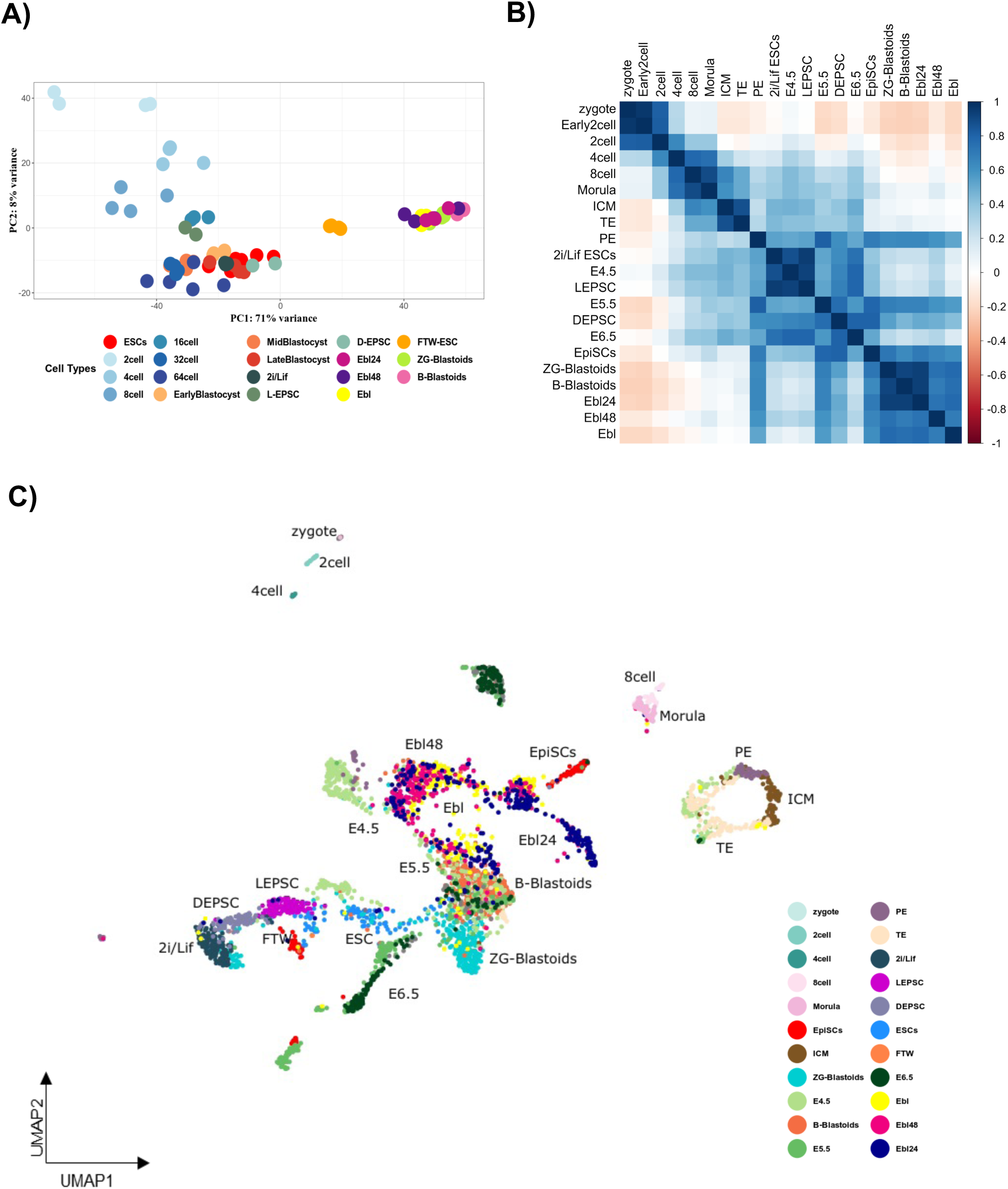
Gene expression analysis of Blastoids using single-cell transcriptomics. A) PCA of sc-RNA seq of E-Blastoids at 24, 48, and 72hrs timepoints after integrating the dataset from this study with published bulk RNA-seq data and sc-RNA-seq data. B) Correlation matrix based on the top 5,000 expressed genes averaged tracking zygote to E6.75, including TE, PE, L-EPSCs, D-EPSCs, 2i/Lif ESCs, EpiSCs, B-blastoids, ZG-blastoids and sc-RNA Seq dataset of Ebl24, Ebl48, and E-blastoid (Ebl). C) Single-cell UMAP analysis after comparing developmental progression from zygote to E6.5. Data sets of sc-RNA seq of Ebl24, Ebl48, Ebl, and blastoids generated by other groups (ZG- and B-blastoids) have been integrated into the same.

**Figure. 5:**
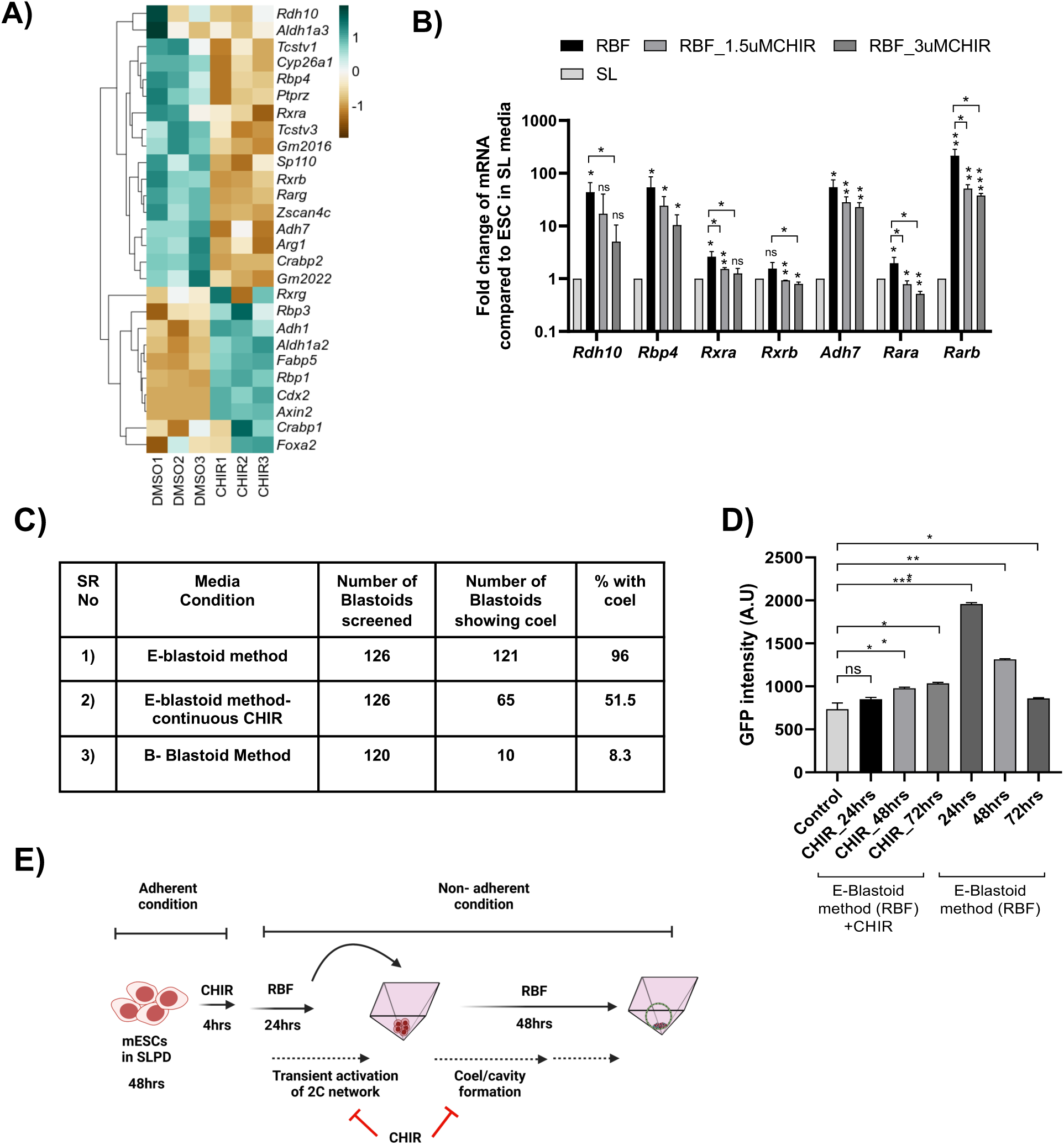
Transient activation of 2C genes is essential for efficient E-blastoid generation from ESCs. (A) Heatmap depicting the expression of different RA-responsive genes in ESCs treated with DMSO and CHIR from the available dataset (GSE40959) (Wu et. al. 2013). Z score blue to red as +1 to -1. (B) Quantitative expression analysis of RA responsive genes in indicated culture conditions, error bar represents SD of the mean of biological replicates (n=3). ‘*’ indicates a p-value <0.05, ‘**’ indicates a p-value <0.01, ‘***’ indicates a p-value <0.001, ‘ns’ indicates a p-value > 0.05. (C) Table showing the percentage of the blastoids with the cavity formed under different conditions. (D) Expression of 2C:GFP reporter in E-blastoid method with continuous CHIR treatment and E-blastoid method at 24, 48, and 72hrs. The error bar represents the SD of the mean of biological replicates (n=3). ‘*’ indicates a p-value <0.05, ‘**’ indicates a p-value <0.01, ‘***’ indicates a p-value <0.001, ‘ns’ indicates a p-value > 0.05. (E) Schematic depicting the stage-specific inhibitory function of CHIR during E-blastoid organization.

To resolve distinct cell type identities during the self-organization of E-blastoids, we performed a comparative analysis scRNA-Seq of the different stages E-blastoid at 24hrs intervals with published scRNA-Seq data of other blastoids and developmental stages(Posfai et al., 2021) (Figure 3C). The E-blastoid cells clustered with cells of E4.5, E5.5, E6.5, B- and ZG-blastoids. This cluster was distinct from the pre-implantation embryos. A majority of Ebl48 and E-blastoid cells clustered between E4.5 and EpiSCs. However, the B-blastoid and ZG-blastoid cells clustered with a different subset of E4.5 cells which closely overlapped with E5.5 and E6.5 (Figure 3C).

Our data shows that all stages of E-blastoids are closely related to other blastoids (B and ZG) and the peri to post-implantation stages of mice embryos from E4.5 to E6.5. All three blastoids (B, ZG, and E) are composed of cells with transcriptomes resembling post-implantation EPI and PE and very little TE lineage. In the hierarchical clustering, all stages of E-blastoids cluster relatively close to E4.5 to E6.5 transcriptome than the B and ZG blastoids suggesting the E-blastoids are relatively closer to the natural embryos (Figure S3A). Collectively, our data suggests that all blastoids, despite their morphological similarities to blastocyst, at best resemble the peri- or post-implantation stages of the embryo at the molecular level, irrespective of the method or initial cell types used for their self-organization.

### Pre-culture of ESCs in SLPD and induction of 2C, TE, PE gene networks result in efficient E-blastoid generation

The efficiency of blastoid generation by multiple methods using mouse stem cells ranges between 12%-36%(Li et al., 2019; Rivron et al., 2018; Sozen et al., 2019; Vrij et al., 2019). Our E-blastoid method achieved around 90% efficiency using ESCs alone, which is comparable to blastoid generations efficiency achieved using naïve hPSCs(Kagawa et al., 2022; Yanagida et al., 2021). The efficiency E-blastoid method can be attributed to two steps in the culture regime. First the adherent pre-culture of ESCs in the SLPD conditions and followed by a suspension culture in RBF (Figure 1A). The ESCs pre-cultured in SL or SLCHIR give rise to some blastocyst like structures at very low frequency suggesting that RBF in suspension culture can induce blastocyst like structures from the ESCs. However, the pre-culture with SLPD sharply increase the efficiency (Figure S1C). Expression of trophoectoderm transcription factors like TFAP2C are suggested to promote the trophoectoderm and blastoid generation potential in human PSCs(Guo et al., 2021; Zijlmans et al., 2022). We analyzed the expression of the trophoectoderm linage genes in SLPD. Although *Tfap2c* and *Cdx2* were not significantly induced, other trophoectoderm genes *Gata3, Tead4, Elf5 and hand1* were significantly induced in ESCs pre-cultured in SLPD (Figure 4A). We suggest the induction of some of the trophoectoderm genes might facilitate the formation of trophoectoderm lineage during the E-blastoid generation in RBF media, and hence pre-culture of ESCs in SLPD is essential for efficient E-blastoid generation.

**Figure. 4:**
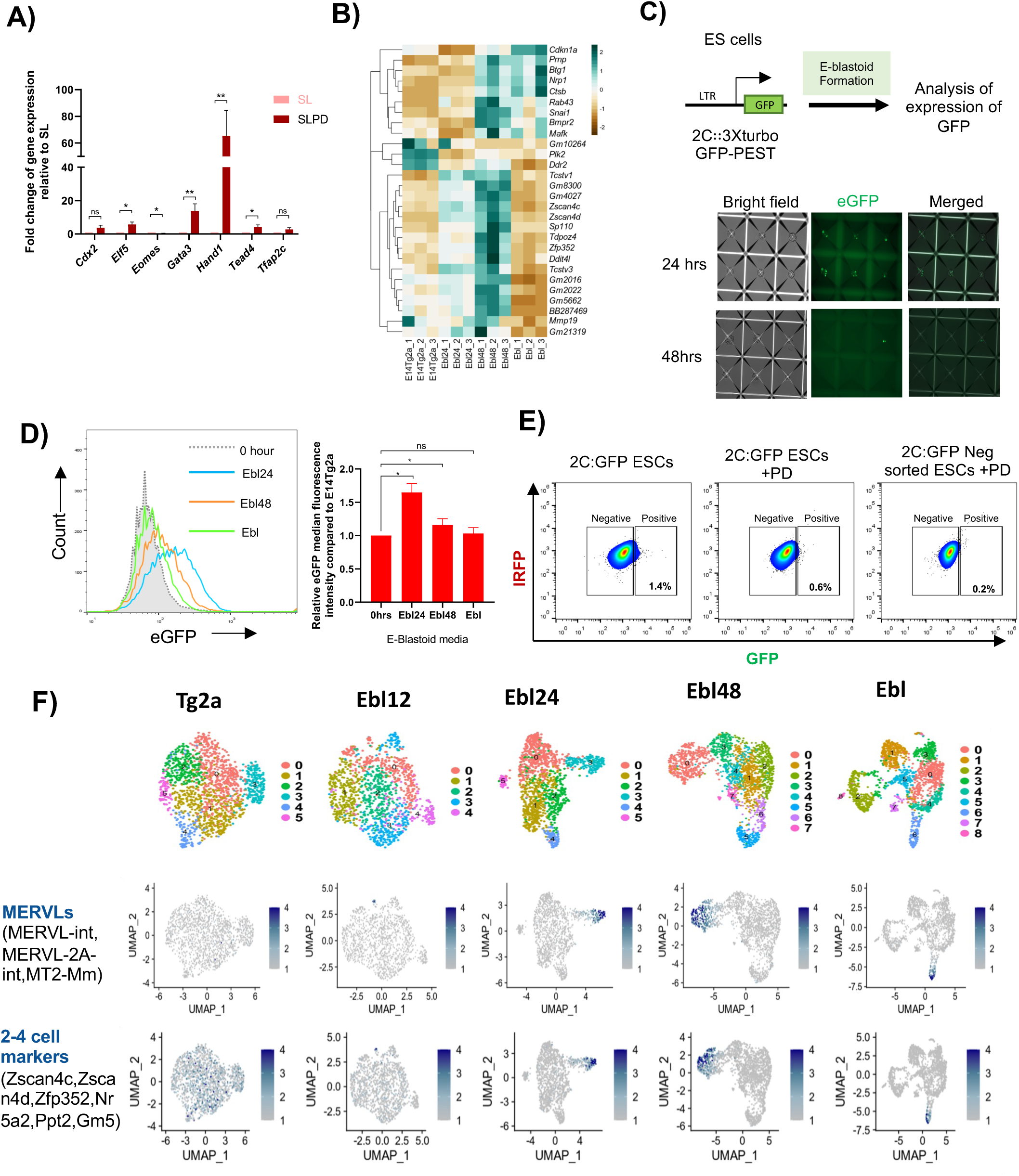
Transient activation of 2C genes is essential for efficient E-blastoid generation. (A) Fold change of gene expression in SL and SLPD. (B) Heatmap depicting the differentially expressed totipotent marker genes in ESCs, Ebl24, Ebl48, and Ebl. Z score blue to red as +3 to -3. (C) (Top) schematic representation of stable 2C::GFP3XturboGFP-PEST reporter in E14Tg2a cell line used for 2C:GFP expression analysis during E-blastoid generation. (Bottom) images showing 2C:GFP expression in E-blastoids during their generation in the Aggrewell at indicated time points. (D) (Left) GFP expression during generation of E-blastoids from 2C:GFP ESC line at indicated time points. (Right) bar plot of the median fluorescence intensity of the GFP expression in 2C:GFP reporter cell line during generation of E-blastoids from 2C:GFP ESC line at indicated time points. The error bar represents the SD of the mean of biological replicates (n=3). ‘*’ indicates a p-value <0.05, ‘ns’ indicates a p-value > 0.05 (E) FACS profile of 2C:GFP expressing cells under depicted conditions. (F) (Top) Single-cell UMAP projection of Ebl12, Ebl24, Ebl48, and Ebl. (Bottom) Feature plots projecting expression of representative MERVLs and 2-4 cell markers overlaying in UMAP.

To further understand, the underlying mechanisms of higher efficiency and the developmental pathways followed by the cells during E-blastoid formation, we analyzed the transcriptome of the E-blastoids at 24hrs (Ebl24), 48hrs (Ebl48) and 72hrs (Ebl) (Figure 4B, S4A). The expression of pluripotency genes and many primed genes (mesoderm and ectoderm) was reduced by 24hrs. The TE and PE genes were induced by 48hrs. While the expression of mesoderm genes was significantly reduced in Ebls, most of the PE and TE genes continued to express (Figure 4B, S4A). Intriguingly the expression of many of the 2-cell-stage (2C) transcripts were induced by 24hrs and continued to express till 48hrs. The majority of 2C transcripts further reduced by 72hrs in Ebls (Figure 4B). MERVL is an LTR-retrotransposon activated in the 2-cell-stage embryo or 2C-like cells (2CLCs) in ESC culture. We generated an ESC line carrying a 2C:GFP-PEST reporter cassette to mark the 2CLCs(Iturbide et al., 2021). The 2C:GFP was actively transcribed in most of the Ebls by 24hrs during Ebl generation (Figure 4C) and decreased by 72 hrs. The Flowcytometry analysis suggested showed a robust induction of 2C: in a large population of cells (13%) in EBLs at 24 hrs, which decreased slightly by 48hrs (3%), and became insignificant by 72hrs (0.8%) (Figure 4D).

The metastable ESCs spontaneously give rise to a small population of 2C like cells which are totipotent in nature. Such totipotent like stem cell states (TBLCs, TLSCs and TPSC) can self organise into blastoids(Xu et al., 2022; Yang et al., 2022; Zhang et al., 2022). There is possibility that E-blastoids may be generated by cells which are already in a totipotent like state and not from the pluripotent population. The preculture in SLPD is essential for high efficiency of E-blastoid self-organization. We cultured the 2C:GFP ESCs in SLPD and analyzed for GFP expression. The GFP expression was significantly reduced in SLPD suggesting that the SLPD treatment reduced 2CLC population the culture (Figure 4E, S4B). Further to eliminate the preexisting 2CLC population, we sorted the GFP negative cells from 2C:GFP cells and cultured them in SLPD. These cells failed to induce GFP expression and could self-organize into blastoids (Figure 4E, S4B). Together, our data demonstrate that the E-blastoids are derived from the non-2CLC population of the metastable ESCs.

We performed single cell RNAseq (scRNA-Seq) at 12-24 hrs intervals to analyze the celluar identities of the cells at different stages of the E-blastoid generation. Consistent with the flowcytometry data, the scRNA-Seq data revealed that MERVL and 2C genes were induced in a significant subpopulation of cells at an early stage (Ebl24) of Ebl generation. This reduced in Ebl48 and was negligible in the Ebls (Figure 4F). The reactivation of the 2C gene network and its subsequent loss during the E-blastoid generation is reminiscent of their expression during embryogenesis between the 2C stage and the blastocyst. The expression of TE genes was prominent in a significant proportion of cells at 48hrs, and the Ebls (Figure S4C). PE genes expression was detected in a distinct subpopulation of cells at 48 hrs, which further increased by 72hrs. The cells expressing pre-implantation-epiblast genes decreased during Ebl generation, with a small proportion of cells expressing them in the Ebls. However, Ebls had a significant cell population expressing the post-implantation-epiblast gene sets (Figure S4C). This is consistent with our immunofluorescence assay data of the E-blastoids that showed that Ebls are composed all the embryonic and extraembryonic lineages of the blastocyst. A higher proportion of post-implantation epiblast-like cells relative to pre-implantation epiblast-like cells in Ebls further strengthens our suggestion that blastoids are mostly similar to peri- or post-implantation stage embryos.

### Transient activation of 2C genes is essential for efficient E-blastoid generation from ESCs

The 2C gene network is expressed after the first cleavage and gradually wanes as the blastomeres specialize to TE and ICM in the blastocyst. The 2C genes are mostly repressed in the EPI and ESC(Falco et al., 2007; Macfarlan et al., 2012). The MERVL activation induces a 2C like state in the ESCs by induction of the 2C genes(Macfarlan et al., 2012; Yang et al., 2020). The 2C genes are the first zygotic genes to be activated in embryos, and they regulate further development of the embryo. The reactivation of 2C genes in significant cell populations during Ebl generation prompted us to investigate if the reactivation of 2C genes in ESCs is essential to efficiently self-organize to blastoids. We analysed the reactivation of 2C:GFP in individual Ebls at 24hrs during their generation, around 63% of them showed 2C:GFP reactivation (Figure S5A). This suggested a strong correlation (r=0.9) between the reactivation of 2C genes and the blastoid generation efficiency.

Our small molecules screen experiments identified RA as an essential component for efficient blastoid generation, blastoid generation from ESC is compromised in absence of RA (Figure S1A). Retinoid signalling induced by RA was shown to reactivate the 2C genes in ESC and efficiently induce 2CLCs(Iturbide et al., 2021). We hypothesized that the RA mediated reactivation of 2C genes during E-blastoids generation might lead to the recapitulation of some of the post-cleavage developmental processes to induce cavitation and TE formation to organize the blastoids.

To test this, we perturbed the RA mediated 2C gene network activation during E-blastoids generation. GSK3β can regulate retinoid signaling in certain cancers(Gupta et al., 2012; Zhang et al., 2020). We asked if GSk3β regulates 2C gene network through retinoid signaling during E-blastoid organization. We analyzed the RNA expression analysis from Wu et. al.(Wu et al., 2013) to evaluate the effect of GSK3β on retinoid signaling in ESCs. The inhibition of GSK3β led to the downregulation of retinoic acid-responsive and retinoic acid pathway genes in ESCs (Figure 5A). This suggested that the GSK3β activity was essential for retinoid signaling and its inhibition by CHIR down regulated retinoid pathway targets. To further ascertain the GSK3β mediated regulation of retinoid signaling, we analyzed the expression of retinoid signaling pathways target genes at 24 hrs of E-blastoids generation, in presence and absence of CHIR. Consistent with scRNA seq data, the retinoic acid pathway genes were induced robustly in the RBF media within 24 hrs. However, the addition of CHIR significantly reduced their induction, suggesting GSk3β promotes RA induced retinoid signaling in ESC during E-blastoid generation (Figure 5B). The incomplete suppression of retinoid pathway genes by CHIR in RBF indicates that the Retinoid pathway in ESC is partially dependent on GSK3β. These data show that GSK3β activity is essential for efficient retinoid signaling and expression of 2C genes in ESCs, suggesting induction of 2C genes by GSK3β-retinoid signaling plays a vital role in efficient E-blastoid generation. This is further supported by the RNA expression data analyzed from Wu et al., where a concomitant downregulation of 2C genes is observed along with down regulation of retinoid pathway (Figure 5A).

To test if GSK3β is essential for efficient E-blastoid self-organization, we supplemented the RBF media with CHIR in the beginning of the E-blastoid suspension culture (Figure S5B). The E-blastoid generation showed an efficiency of 96% which was drastically reduced to 52% in presence of CHIR (Figure 5C). We further analyzed the expression of 2C:GFP during E-blastoid generation. The addition of CHIR severely impaired the induction of the 2C:GFP expression, its expression was barely detectable at 24 hrs in contrast to robust induction in absence of CHIR. A marginal induction of GFP was seen at 48 and 72hrs, suggesting GSK3β is essential for the transient reactivation of 2C genes during E-blastoid generation (Figure 5D). The induction of 2C:GFP by 24hrs followed by its downregulation during the later stages of E-blastoid generation despite the presence of RA suggests that the 2C genes respond to RA and activate retinoid signaling and 2C genes only in the early stages of E-blastoid organization (Figure 5D).

Different methods of generation of blastoids from mouse ESCs and TSCs have used CHIR in their culture media with 12-36% efficiency(Vrij et al., 2019). E-blastoid generated in our study and B-blastoids generated by Li et. al. has not used TSCs for blastoid generation, necessitating the TE compartment contribution from the pluripotent stem cells. The method used by Li et al to generate B-blastoids from EPSC uses a media containing CHIR(Li et al., 2019). We also used this method to assess the role of CHIR on 2C gene network induction (Figure S5B). We cultured the 2C:GFP cells in EPSC conditions and generated B-blastoids as described in Li et. al. In our hands, the B-blastoid efficiency was around 8%, and the E-blastoids at 96% (Figure 5C). Unlike robust induction of 2C:GFP in E-blastoids by 24hrs (Fig 5D), the median of 2C:GFP expression was significantly reduced until 48hrs in the B-blastoid method only to recover back to levels comparable to the ESC by 72hrs (Figure S5C). Further, The 2C:GFP was induced in very few blastoids (3.4%) during B-blastoids generation compared to 62% during E-blastoids (Figure S5D). These results suggest that GSK3β inhibitor represses 2C gene expression in the early stages of B-blastoid generation much below the levels in ESC. We suggest that this could one of the major reasons for poor efficiency in the B-blastoid method.

Collectively our data shows that, RA transiently induces the 2C gene network in ESCs in the early stages of E-blastoid generation. GSK3β is essential for efficient induction of 2C gene network in ESCs and the early stages of blastoid generation. Together our data demonstrate that transient activation of 2C genes by GSk3β-retinoid signaling during blastoid generation enhances the efficiency of cavitation and blastoid generation from ESC.

## Discussion

Here, we present evidence of self-organization mESC into blastoids at high efficiency. The morphology, cell lineage composition, and transcriptome profile of the E-blastoids are similar to the blastocysts. In contrast to current understanding, our data show that the pluripotent mESCs possess full potential to self-assemble into blastoids composed of all three blastocyst lineages. The mESC attain this potential by induction of some of the trophoectoderm genes *Gata3*, *Tead4*, *Hand1* and *Elf5* in pre-culture in SLPD and by transient reactivation of totipotency gene network in the early stages of E-blastoids. The underlying mechanisms of dependence on GSK3β activity in transient reactivation of totipotent gene network during E-blastoid generation by retinoid signaling would require further investigation.

All blastoids are composed of all three lineages of balstocyst, the efficiency of generating them depends on the availability of each of these cell types(Vrij et al., 2019). It remains unclear whether blastoid generation from constituent stem cell types, such as ESCs and TSCs, follows the transcriptome and developmental sequence of preimplantation embryogenesis(Rivron et al., 2018). TBLCs, TLSCs, and TPSCs have transcriptome and developmental potential similar to totipotent blastomeres. Therefore, the blastoid generation process from these cells is expected to follow a molecular and developmental course similar to the cleavage to blastocyst. mESCs resemble E4.5 epiblast, established after the segregation of the two extraembryonic layers. However, the induction of trophoectoderm genes in SLPD and co-existence of distinct population with 2C transcriptome and TE transcriptome in EBL48 suggest two complementary paths may be simultaneously operating to generate E-blastoids efficiently. One by differentiation of some of the ESC population into TE lineage, and other by reactivation of 2C network in subpopulation of ESCs leading to recapitulation of the developmental sequences from cleavage to blastocyst. The possibility of differentiation of some of the ESCs to TE is strongly supported by our recent report where we have shown that mESC and the epiblast possess potential to differentiate trophoectoderm.

Human PSCs are equivalent to the human blastocyst’s epiblast and can readily differentiate to trophoectoderm(Guo et al., 2021). 8-cell like cells (8CLCs) with a totipotent gene network can arise in the hPSC culture in a manner similar to totipotent 2CLCs in mESCs(Mazid et al., 2022; Taubenschmid-Stowers et al., 2022). Blastoid generation from hPSCs has been reported to follow some developmental sequences, however the reactivation of the totipotency or 8CLC gene network has not been analyzed at early stage of blastoid generation. The capability to differentiate into trophoectoderm, the capacity to produce transitory populations of totipotent-like cells(Macfarlan et al., 2012; Mazid et al., 2022), and the sharing of a comparable hPTMS(Zijlmans et al., 2022) characterize the naive hPSCs and mESC. We suggest that the naïve hPSCs, mESCs and PSCs of other mammals possess similar basic developmental potential. The differences observed in molecular pathways governing various mammalian PSCs could be responsible for the variations in the difficulty/routes in realizing their full development potential.

All the blastoids reported till date can develop in some form till mid gestation. The current focus is to develop methods to develop blastoids to achieve later developmental stages. The cellular composition and the quality of the blastocyst is known to impact the implantation and post-implantation development in mice and humans(Bolton et al., 2016; Gardner et al., 2000). We suggest that the understanding the mechanisms of blastoid generation which is not given as much attention will be critical to develop better quality blastoids to attain latter phase embryonic development.

## Acknowledgements

We thank Mr. Jedy Jose for technical support in embryology experiments. D.J, V.V.V, were supported by a fellowship from UGC (India). H.T.K was supported by a fellowship from ICMR (India). We thank the Microscopy, FACS, Animal House and transgenic core facilities of CCMB for the support extended to carry out this work. P.Si acknowledges the stipend support from the DBT grant GAP0546. WT/DBT India Alliance grant 500053/Z/09/Z -P.C.S., Department of Biotechnology grant BT/PR14064/GET/119/16/2015 -P.C.S., DBT grant GAP0582: BT/PR40264/BTIS/137/44/2022 -D.T.S.

## Author contributions

Conceptualization, D.J, and P.C.S Methodology, D.J, P.Si, P.Sa, N.K, A.K, J.L, and P.C.S; Investigation, D.J, P.Si, P.Sa, and D.T.S; Writing – Original Draft, D.J, P.Si and P.C.S.; Writing – Review & Editing, D.J, P.Si, D.T.S and P.C.S.; Funding Acquisition, D.T.S and P.C.S.; Resources, D.J, V.V.V, and H.T.K; Visualization, P.Si and D.J.; Supervision, P.C.S.

## Methods

### Culture of metastable and ground-state mouse ES cells

Metastable ES cells were cultured on 0.1% (w/v) gelatine-coated tissue culture-treated dishes in serum + LIF (SL) media containing 10% (v/v) heat-inactivated FBS in GMEM (12.5g/litre w/v), NaHCO3 (32.7mM), Sodium Pyruvate (1mM), NEAA(0.1mM), β-mercaptoethanol (0.1mM) and 1000U/ml LIF. The cells were grown in a humidified incubator under 5% CO2 at 370C. Ground state ES cells were cultured on 0.1% gelatine coated tissue culture-treated dishes in N2B27, containing DMEM/F12 and Neurobasal medium in 1:1 (v/v) ratio, supplemented with N2 supplement (1X), B27 supplement (1X), NaHCO3 (32.7mM), Sodium Pyruvate (1mM), NEAA (0.1mM), and β-mercaptoethanol (0.1mM) along with recombinant hLIF (1000U/ml), CHIR99021 (3μM) and PD0325901 (1μM). Upon 70% confluency, TrypLE-EDTA was used for dissociation and passaging.

### Extended pluripotent stem cell culture

Extended pluripotent stem cells (EPSCs) were converted from metastable ES cells when grown in culture media as described by Hongkui Deng’s group (Yang, et al. 2017) where basal N2B27 media was supplemented with small molecules and cytokines as follows: 10 ng/ml recombinant hLIF, CHIR99021 (3µM), (S)-(+)-Dimethindenemaleate (2µM), Minocycline hydrochloride (2µM) and 5mg/ml BSA. Upon 70% confluency, EPSCs were passaged with TrypE-EDTA, and splitting was done in a 1:3 ratio.

### Generation of E-blastoids from ESCs and B-blastoids from EPSCs

ESCs were seeded on a 0.1% gelatine-coated tissue culture dish and cultured in SL media supplemented with PD0325901 (final 1μM) for 48 hours. The cells were trypsinized using TrypLE-EDTA and neutralized with SL media (without LIF). Around 48,000 cells were seeded per well of 24 wells of Aggrewell™400 containing around 1ml of RBF media and centrifuged at 100 rcf for 3 minutes. Aggrewell™400 was prepared by rinsing the wells with an anti-adherence rinsing solution (Stem cell technologies) as per the manufacturer’s protocol and washed with SL (without LIF) before seeding the cells. RBF media was composed of GMEM (12.5g/litre) with 10% FBS (v/v), NaHCO3 (32.7mM), Sodium Pyruvate (1mM), NEAA (0.1mM), β-mercaptoethanol (0.1mM) supplemented with Retinoic acid (1µM), bFGF (25ng/ml) and BMP4 (20ng/ml). 24 hours later, 500µl of RBF was added carefully without disturbing the cell aggregates. After 48hrs, cavitation was observed in most of the aggregates. CHIR (3µM) was added to RBF at this point. Blastocyst-like structures were observed in most of the Aggrewells by 72 hrs. EPSC was derived from the ESC as described by Yang et. Al. (Yang et al., 2017a). B-blastoids were generated from the EPSCs as described by Li et al.(Li et al., 2019).

### Derivation of ES-, TS- and XEN-like cells from blastoids

For the derivation of TS-like cells from the E-blastoids, we followed the method described by Ohinata et. al.(Ohinata and Tsukiyama, 2014). For the derivation of ES- and XEN-like cells, single blastoids were seeded onto feeders cells in 96 wells. Blastoids were attached within 2-3 days onto feeders and outgrowth was observed. Outgrowth was dissociated using TrypLE-EDTA and the cells replated onto freshly seeded feeders. For ES-like cell derivation, the cells from outgrowth were cultured in N2B27 media supplemented with hLIF (1000U/ml), CHIR99021 (3μM), and PD0325901 (1μM). For XEN-like cell derivation, a protocol by Rugg-Gunn et al. (Rugg-Gunn, 2017) was followed. The cells from outgrowth were seeded onto feeders and cultured in TS medium +FGF4. TS medium constituted of RPMI 1640, FBS (20% v/v), Sodium pyruvate (1 mM), β-mercaptoethanol (100 μM), L-glutamine (2mM) supplemented with 25ng/ml of FGF4 and heparin (1µg/ml). the XEN-like cells were passaged at 80% confluency. Finally, XEN-like cells were cultured on a 0.1% gelatine coated tissue culture dish with a 70% feeder conditioned media and 30% TS medium.

### Mice

C57BL/6J and CD1 mice were obtained from Jackson laboratories. All the procedures related to mice models were performed following the ethical guidelines of the Institutional Animal Ethics Committee (IAEC) of CCMB and approved by the Committee for Control and Supervision of Experiments on Animals (CPCSEA) before any experiments. For pseudopregnancy, CD1 female mice of 8 – 11 weeks old in the estrus cycle were mated with vasectomized F1 male mice. All the mice were housed in a 12hr light/ 12hr dark cycle in a controlled temperature (18-220C) and humidity (R.H. 40-70%) facility and also provided with free access to water and food.

### Blastoid and blastocyst transfer for implantation

E-blastoids, B-blastoids, and blastocysts were washed thrice in the M2 medium. 40 blastoids were transferred into single pseudo-pregnant CD1 mice. A C-section was performed at 6.5 dpc. For visualizing the implantation sites, 0.1ml of 1% Pontamine sky blue was injected through the tail vein using a 27-gauge needle 10 minutes before sacrificing the mice. Deciduae were dissected out of the uterus and fixed with 4% PFA overnight for immunohistochemistry.

### Generation of chimeric mouse from Ebl-derived ES cell line

1.5dpc mouse embryos were collected from C57BL/6J mice. 4-6 cells of Ebl-derived ES cell line were microinjected into blastocyst stage of the mouse embryos obtained from C57BL/6J mice. 18 microinjected blastocysts were transferred to single pseudo-pregnant CD1 mice. Pups born were checked for coat colour for chimerism.

### Histology of E6.5dpc mouse embryo and implanted E-blastoid

Deciduas at E6.5 were dissected from the CD1mouse uterus and immediately fixed with 4% paraformaldehyde and kept overnight at 4oC. Fixed tissues were dehydrated and embedded in wax. Sections of 8-micron thickness were taken on ProbeOn slides and followed for Hand E staining.

### Immunofluorescence staining

Cells were cultured in 2D culture in 24 wells for up to 70% confluency. Cells were washed thrice with 500µl of PBS and 500 µl of freshly prepared 4% paraformaldehyde (made in PBS) fixative was added to the plate and incubated at RT for 20 minutes. The fixative was removed and the plate was washed thrice with 1ml PBS. The specimen was blocked and permeabilized in a blocking buffer containing PBS with 5% (w/v) BSA, and 0.3% (v/v) Triton-X 100 for 60 minutes at room temperature. Primary Antibodies were diluted in antibody dilution buffer (ADB) containing PBS with 1% (w/v) BSA, and 0.3% (v/v) Triton-X 100. Specimens were incubated with primary antibodies for 4oC overnight. Cells were washed 3 times with PBS and incubated with secondary antibodies diluted at 1:1000 in ADB. Cells were washed with PBS and mounted with Vectashield (H-1200) containing DAPI.

For staining 3D structures like embryos or blastoids, they were first washed twice with PBS. Structures were fixed in 4% paraformaldehyde in PBS for 15 minutes and rinsed in PBS containing 3 mg/ml polyvinylpyrrolidone (PBS/PVP). Thereafter structures were permeabilised in PBS/PVP containing 0.25% Triton X-100 for 30 minutes. Blocked in blocking buffer, comprising PBS containing 5% BSA, 0.01% Tween 20 for 60 minutes. Primary antibodies were diluted with the appropriate antibody dilution as per the manufacturer protocol in PBS containing 1% BSA, and 0.01% Tween 20 and incubated at 4°C overnight. They were rinsed three times in blocking buffer for 5 minutes each and incubated with secondary antibodies diluted as 1:500 in PBS containing 1% BSA, 0.01% Tween 20, and incubated for 60 minutes at room temperature. Rinsed 3 times with PBS and stained for nuclei with DAPI (1µg/ml) prepared in PBS for 15 minutes at RT. Embryos/blastoids were finally rinsed 3 times in PBS and images were acquired by confocal microscopy.

### Real-time PCR analysis

The RNA was extracted from 1 million cells with TRIzol by manual method and quantified by Nanodrop (Thermo Scientific). One microgram of RNA was reversed and transcribed into cDNA by using SuperScript™ III First-Strand Synthesis System. The first strand synthesized cDNA was diluted 5 times and real-time PCR was set with power SYBR Green PCR master mix on the ABI 7900 HT. The PCR setup was as follows: Step 1: 95^0^C for 10 min, step 2: 95°C for 15 sec, step 3: 60°C for 30 sec, and step 4: 72°C for 30-sec steps 2-4 was repeated for 40 cycles. GAPDH was used as an internal control. The reactions were analyzed by the software (SDS 2.1) provided with the instrument. The primers used for real-time PCR are given in the resource table.

### Western blot analysis

The cells were harvested by scraping them from the plates in PBS and collected by centrifugation. The cell pellet was washed twice with PBS and reconstituted in RIPA. RIPA buffer constituting 25mM Tris HCl (pH 8.0),150mM NaCl, 1% NP-40, 0.5% Sodium deoxycholate, 0.1% SDS, and Complete Protease Inhibitor Cocktail Tablets (Roche). The lysate was sonicated in Bioruptor (Diagenode) and centrifuged. The clear supernatant was transferred to the fresh tube. The protein samples were denatured in loading dye containing β-mercaptoethanol and resolved by SDS-PAGE. Resolved samples were then transferred onto a polyvinylidene difluoride (PVDF) membrane and blocked-in blocking buffer containing 5% non-fat milk in TBST. Primary antibody hybridisation was carried out overnight at 4°C in 3% non-fat milk in TBST. After incubation three washes were performed with TBST. The secondary antibody was diluted at 1:10,000 in 3% non-fat milk in TBST and incubated for an hour at RT. After incubation three washes were performed with TBST and visualized using enhanced chemiluminescence (ECL) detection kit (Thermo Scientific) and developed in Chemi doc MP (Biorad).

### FACS analysis

70% confluent culture was made into single-cell suspension using TrypLE for 4 mins at 37oC. The cells were diluted in media and pelleted by spinning at 300g for 5 mins. The media was removed and around 1 million cells were resuspended in 300 µl of PBS containing 2% FBS. The cell samples were directly taken for analysis either in Gallios (Beckman Coulter B5-R1-V2) FACS analyzer or LSR Fortessa (BD) analyzer and data was recorded. The FACS data were analyzed using FlowJo vX.0.7

### Bulk RNA-seq library preparation

Using the TRIzol reagent and the manufacturer’s instructions, total RNA was extracted from ESCs and Ebl at various times. Total RNA of 1μg was used to generate libraries using of Illumina stranded total RNA prep with Ribo-Zero Plus library preparation kit (Illumina, 20040529). The Qubit dsDNA HS (High Sensitivity) Assay Kit (Invitrogen, Q32854) was used to measure library concentration, and different libraries were combined to create the final pool at an equimolar ratio. Using an Illumina NovaSeq 6000 instrument, libraries were sequenced for read length of 151 bp reads and a read depth of approximately 30 million reads.

### Bulk RNA-seq data analysis

Paired-end Bulk RNA sequencing was done for ESCs and E-Blastoids. The quality assessment of raw data was done by FastQC v0.11.9 and Illumina universal adapter content was removed using cutadapt v2.8. The filtered sequencing reads were mapped to mouse reference genome mm10 and Gene counts were obtained by STAR_2.5.4b. The data showed an average of 81% uniquely mapped reads which were annotated with Ensembl database. Transcripts Per Kilobase Million (TPKM) were calculated by the rsem-calculate-expression function of RSEM v1.3.3. DESeq2 v1.30.1 was used to get the differentially expressed genes based on the Bayes theorem. Genes showing expression of at least 50 in a row were retained for further analysis. Principal component analysis (PCA) and heatmap analysis were performed with the functions plotPCA and pheatmap in R. The visualization of differential expression of marker genes was performed on TPM counts after scaling and normalizing the read counts by row.

### Single RNA-seq library preparation

Single cell suspension was made from Ebl12, 24, 48 and 72 using TrypLE-EDTA solution. Live cells were sorted as PI-negative population using FACS sorter. Approximately 5000 cells were loaded for Library preparation using Chromium Next GEM Single Cell 3’ Reagent Kits v3.1 (10x genomics). Libraries were sequenced for read length of 91 bp reads.

### Single-cell RNA-seq data processing

Chromium single-cell RNA-seq Illumina base call files (BCL) for FACS sorted Ebls (12, 24, 48, and 72hrs) and E14Tg2a(ESCs) were demultiplexed by using the mkfastq function of cellranger v6.0.2 which is specific to 10X libraries. A customized reference was made to add the repeats in the reference file by the cellranger mkref pipeline. Reads were aligned against mouse reference mm10 and filtered by the count pipeline to get feature, barcode and gene matrices. The R package Seurat v4.0.1 was used to analyze the feature-barcode matrix. The quality assessment of data was done by scater v1.18.6. Scrublet, a Python tool, was used to calculate doublet scores and predictions. Cells with more than 500 detectable genes with a doublet score of < 0.25 and expression of mitochondrial genes accounted for less than 5% of total expression were filtered from the dataset for further downstream analysis. Normalisation and variance stabilization was done by transform and variations due to genes were regressed out. Cells were clustered with the FindClusters function with a resolution of 0.5 and visualization was done using the RunUMAP function. AddModuleSCore function was used to get the aggregate expression of markers and visualized using Featureplot. The additional raw sequence files and preprocessed count matrices were downloaded as per the availability provided in (Gene Expression Omnibus (GEO): GSE14509, GSE45719, GSE84892, GSE100597, GSE74155, GSE135289, GSE135701, GSE134240, GSE130957, ENA: ERP005641. The SmartSeq2 datasets were processed and Gene counts were calculated by STARaligner same as the Bulk seq data processing.

### Single-cell gene expression analysis of merged datasets

All the datasets were analyzed using Seurat v4.0.1 and DESeq2 v1.30.1 in R version 4.0.5. The data processed from all the studies were downscaled and down-sampled before integration and integrated with the present study by merge function from the Seurat suite. The top 2000 differentially expressed genes for PCA computation were selected by the FindVariable function after log normalization and scaling of single-cell data. The top 15 principal components were used for the calculation of UMAP coordinates. The UMAP embedding consisting of all the pre-implantation lineages was constructed using shared nearest neighbor (SNN) modularity optimization implemented in Seurat’s FindClusters function. Expression of individual Marker genes of specific cell lineage was visualized in the E-blastoids population using FeaturePlot from Seurat. The batch effects among the multiple datasets were removed using limma in R. PCA of the merged log transformed datasets was performed by taking an input of the top 1000 expressed genes using the plotPCA function from DESeq2 with variance stabilizing transformation. A correlation plot based on spearman correlation was produced by providing the log normalized merged count matrix of the top 2000 genes to corrplot v0.88.

### Data and materials availability

All the cell lines and plasmid constructs used in this study will be made available against and email request and Material Transfer Agreement (MTA). Raw and processed transcriptome data is deposited in NCBI GEO and is available under the accession GSE219001. The code used for analysis and visualization of scRNAseq data is available at https://github.com/SowpatiLab/Eblastoids.

### Statistical analysis

Statistical analysis was done by using a two-tailed paired student t-test. The representation of data is in the form of means+/-SDM. The was calculated for more than three independent experiments P value<0.05 is considered statistically significant. * represents P<0.05, ** represents P<0.01, *** represents P<0.001, and **** represents P<0.0001.

## Supplemental information titles and legends

**Fig. S1.**
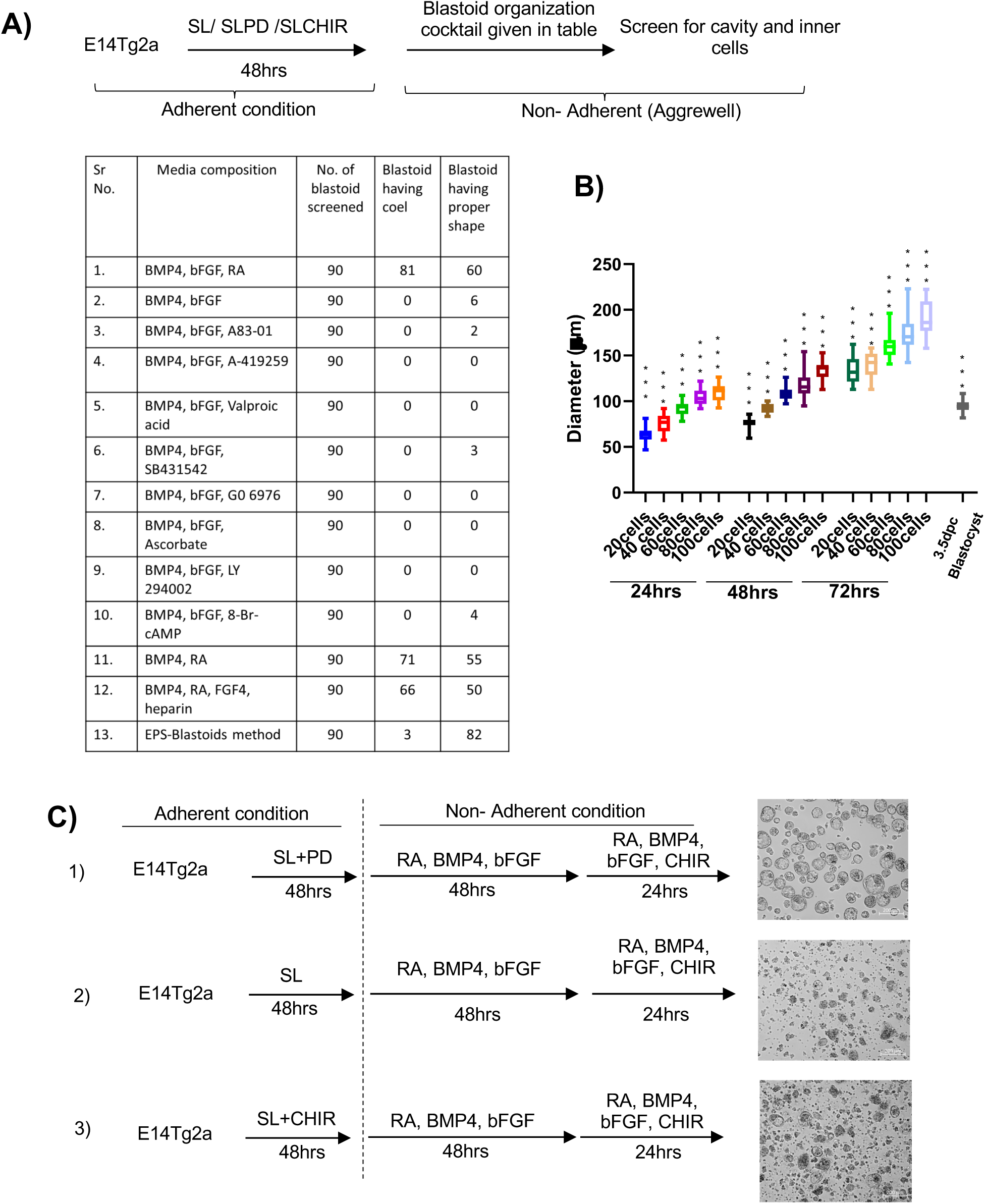
Generation of Blastoids from mESCs, Related to Figure 1. (A) (top) Schematic followed for small scale screen to generate blastoids from ESCs. (Bottom) table enlisting some of the various combinations of small molecules and cytokines used in the screen. (B) The diameter of the E3.5 blastocyst and E-blastoids at 24hrs, 48hrs, and 72hrs with starting cell numbers of 20, 40, 60, 80, and 100 (n=20). (C) (left) Schematic followed for E-blastoid generation in different adherent and non-adherent stages. (right) Phase contrast image of the blastoid generated from these methods (scale bar = 100µm). ‘*’ indicates a p-value <0.05, ‘**’ indicates a p-value <0.01, ‘***’ indicates a p-value <0.001, ‘ns’ indicates a p-value > 0.05.

**Fig. S2.**
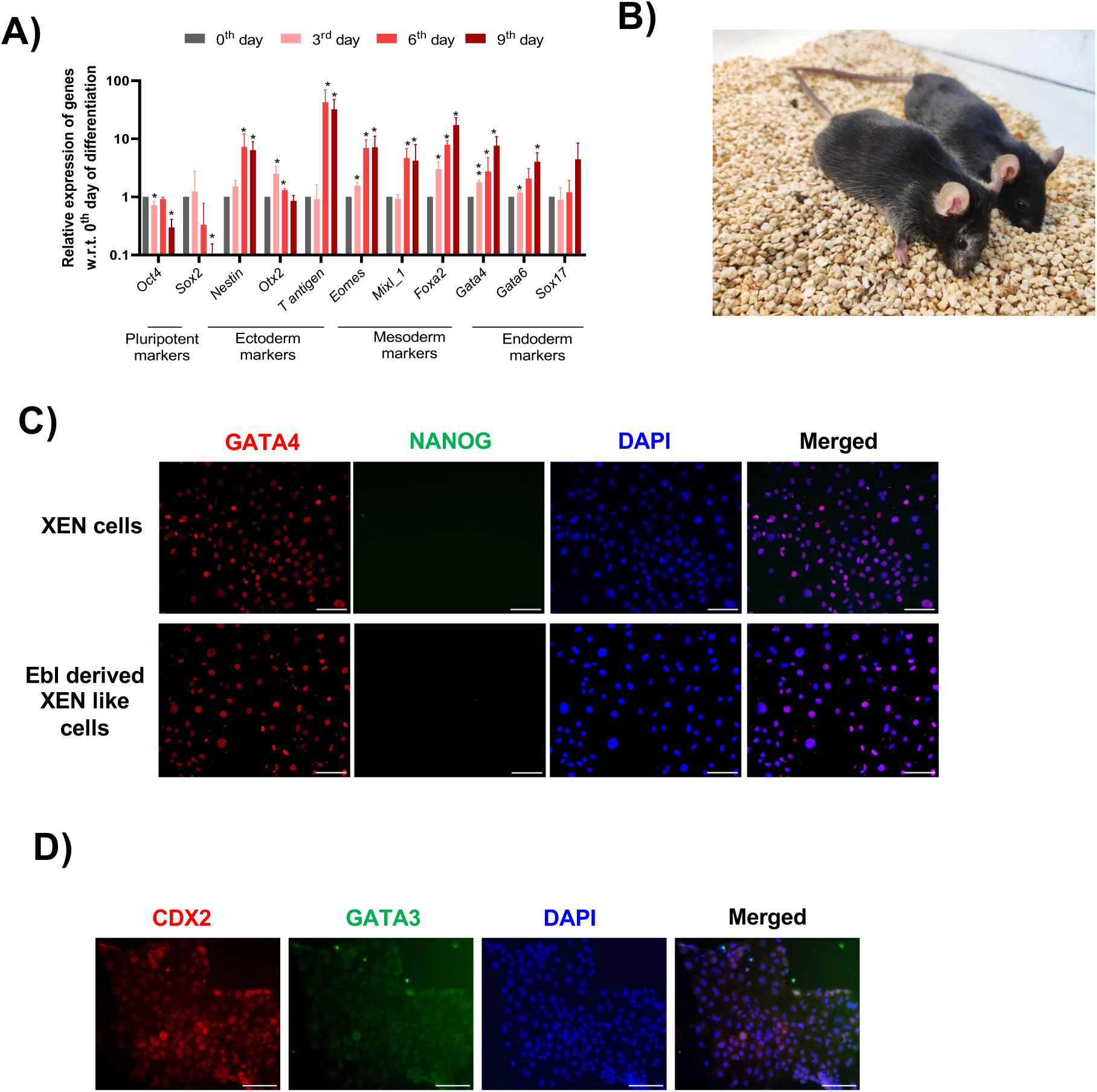
Differentiation potential of E-Blastoid, Related to Figure 2. (A) Relative mRNA abundance of different developmental genes from Ebl-derived ESCs at the 3rd, 6th, and 9th day of embryoid body differentiation analyzed by q-RTPCR. The error bar represents the SD of the mean of biological replicates (n=3). ‘*’ indicates a p-value <0.05. (B) Image of 21-day-old chimeric mice generated from blastocyst injection of ESCs derived from Ebl into C57BL6 blastocyst. (C) Immunofluorescence of GATA4 and NANOG in XEN cells and XEN-like cells derived from Ebls. Scale bar = 100µm. (D) Immunofluorescence of CDX2 and GATA3 in TS-like derived from Ebls. Scale bar = 100µm.

**Fig. S3.**
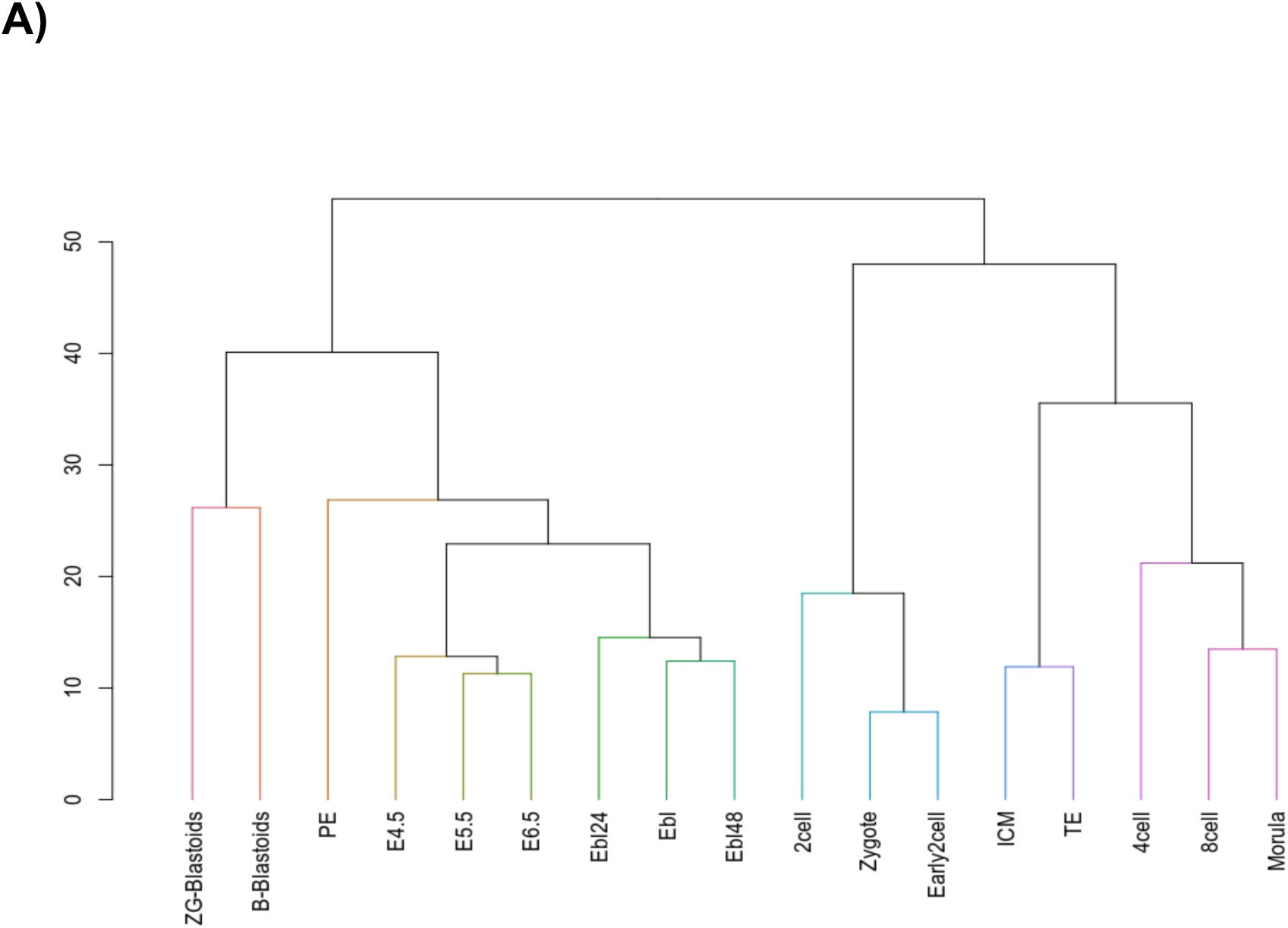
Single-cell Transcriptome analysis of E-blastoids, Related to Figure 3. A) Hierarchical clustering of cell types generated from scRNA seq from Fig 3B.

**Fig. S4.**
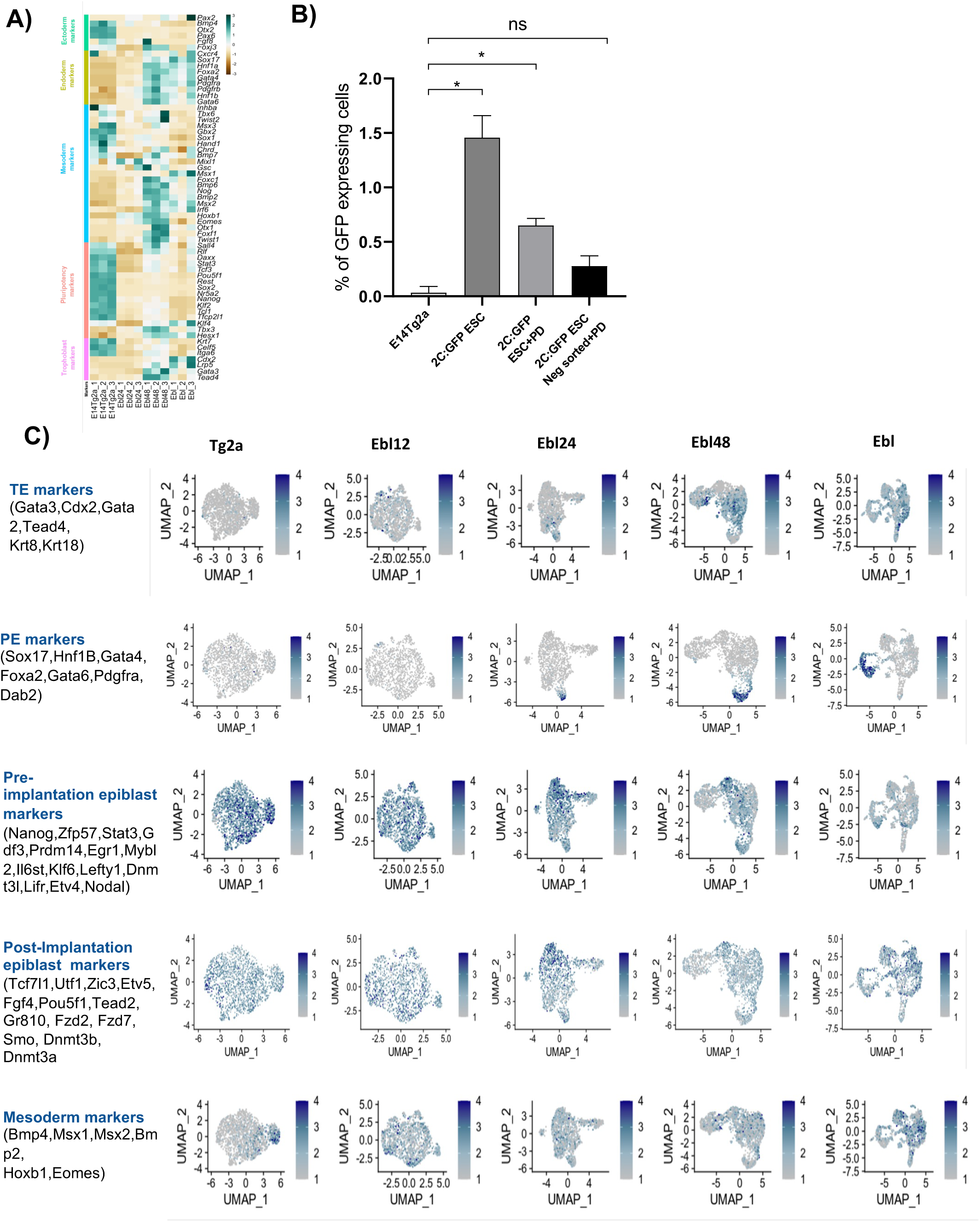
Gene expression analysis in E-blastoids at different time points, Related to Figure 4. (A) Heatmap depicting the differentially expressed genes in ESC, Ebl24, Ebl48, and Ebl. Z score blue to red as +3 to -3. (B) Quantification of the percentage of the GFP-positive cells in Fig 4E. (D) Feature plots projecting expression of representative TE, PE, pre-implantation epiblast, post-implantation epiblast, and mesoderm markers overlaying in Fig 4D UMAP.

**Fig. S5:**
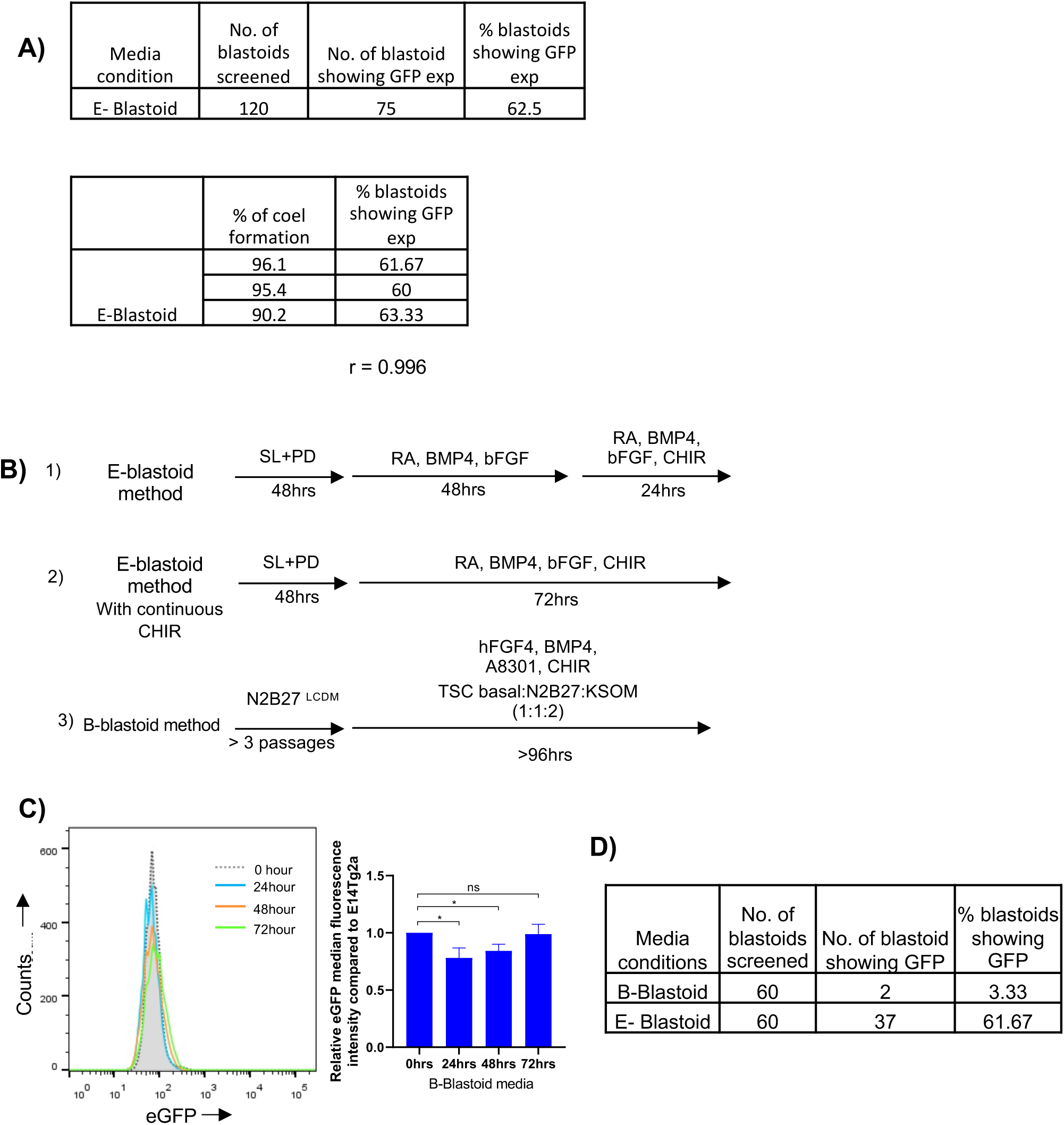
Transient activation of 2C genes is essential for efficient E-blastoid generation from ESCs, Related to Figure 5. (A) (Top) Table showing the percentage of E-blastoids expressing 2C:GFP. (Bottom) Table showing the correlation between cavity formation and 2C:GFP expression during E-blastoid generation. (B) Schematic representing the protocol followed for blastoid formation. ESCs were first grown in SLPD for 48hrs in adherent conditions followed by different culture regimes - 1) E-blastoid method, 2) E-blastoid method with continuous CHIR treatment, 3) E-blastoid method without CHIR 4) E-blastoid method with LIF. (C) (Left) histogram profile of GFP expression in 2C:GFP reporter cell line at 0, 24, and 48hrs time points in B-Blastoid method. (Right) bar plot of the median fluorescence intensity of the 2C:GFP expression at 0, 24, 48, and 72hrs during B-Blastoid generation. The error bar represents the SD of the mean of biological replicates (n=3). ‘*’ indicates a p-value <0.05, ‘**’ indicates a p-value <0.01. (D) Table showing the percentage of blastoids showing 2C:GFP expression in B and E-Blastoid media.

## Key resource table

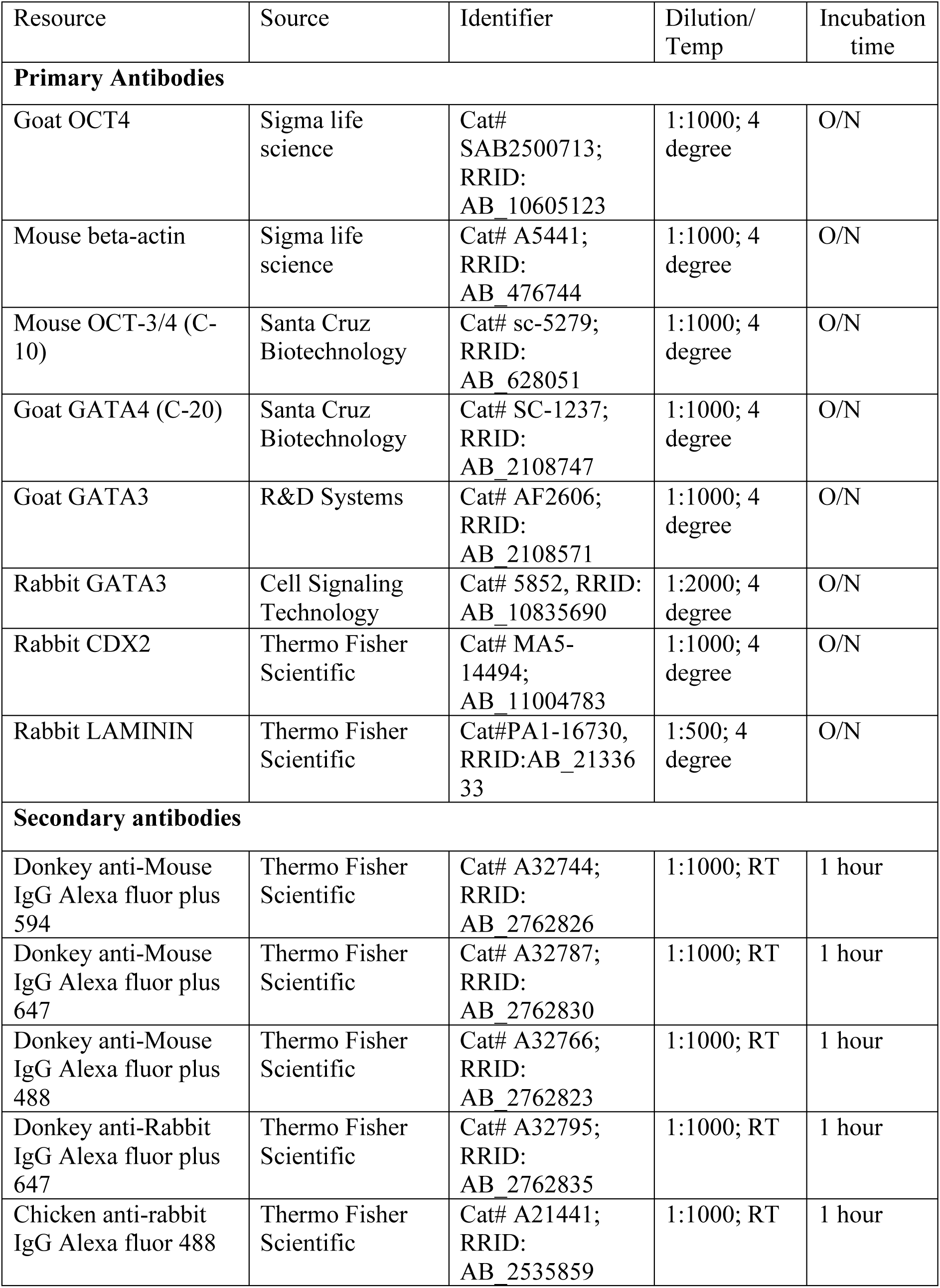
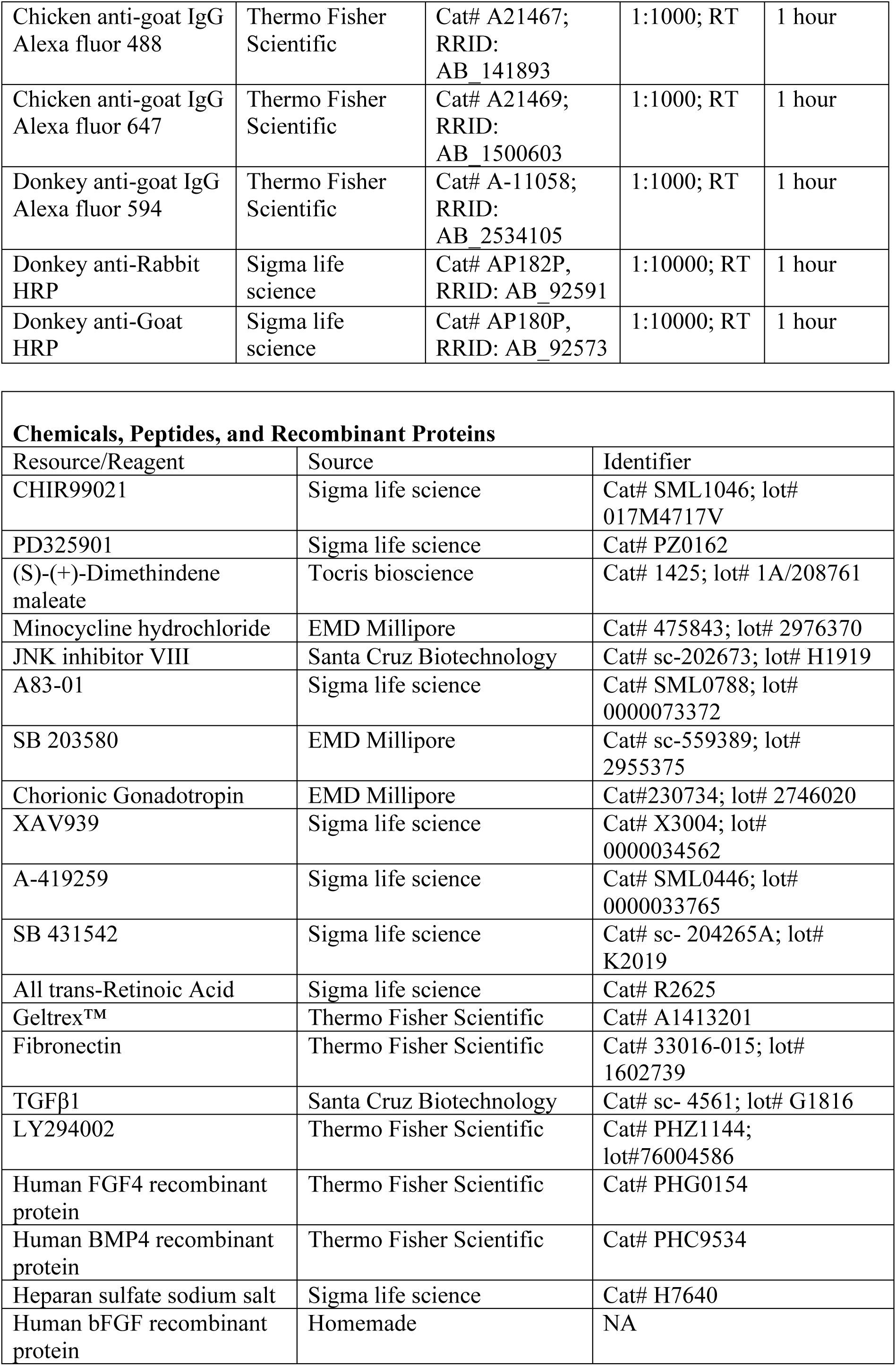
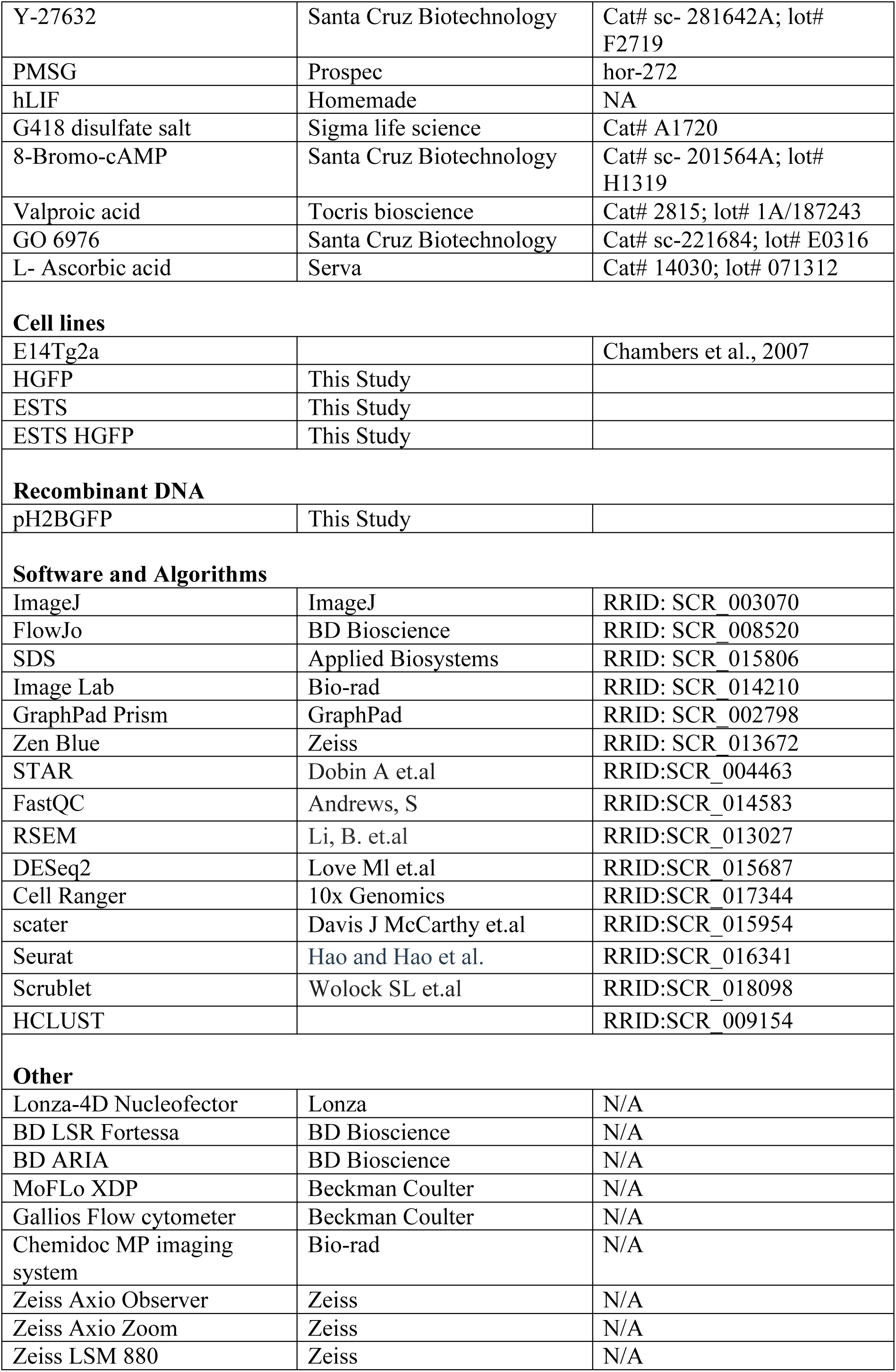
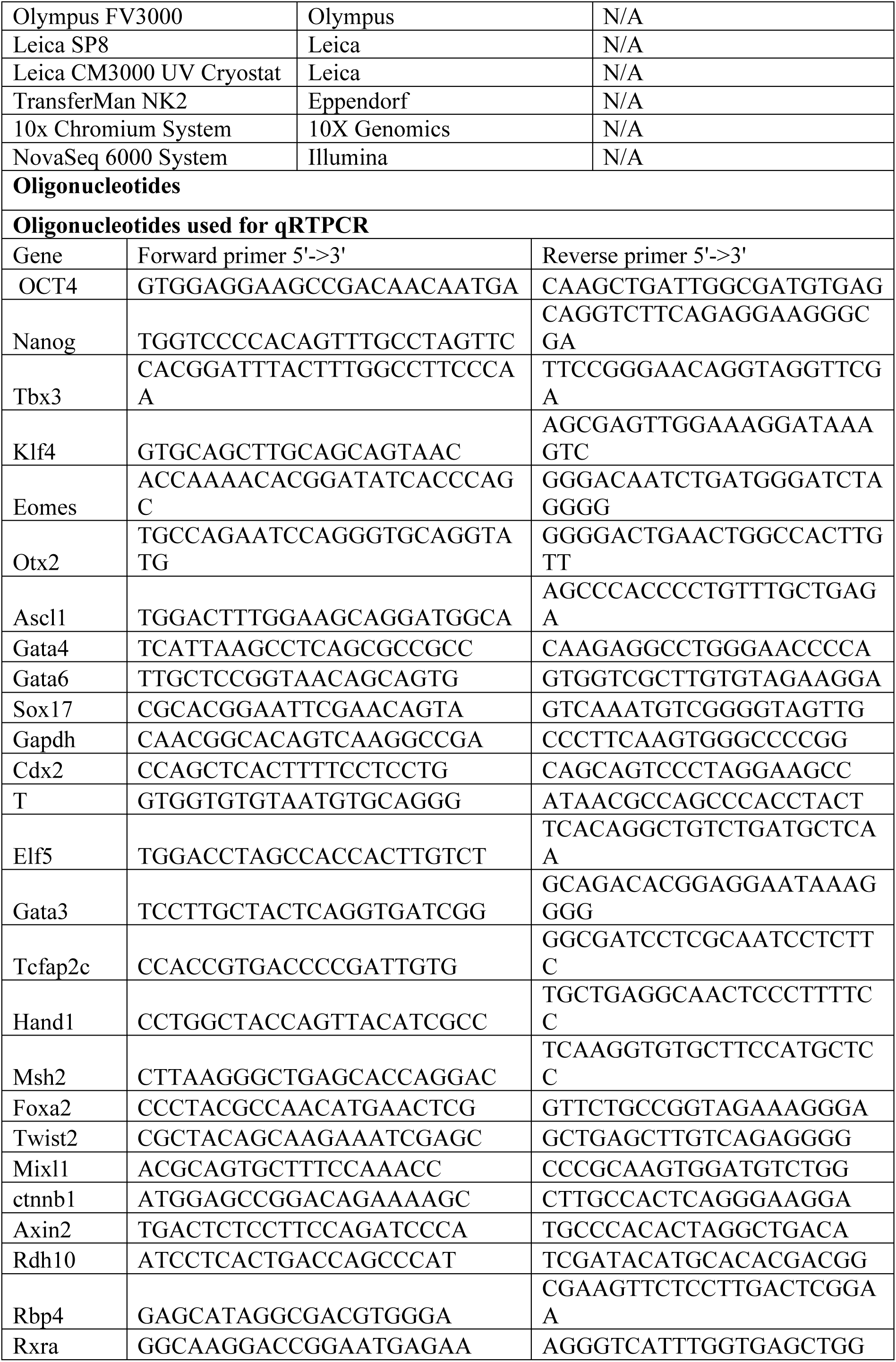
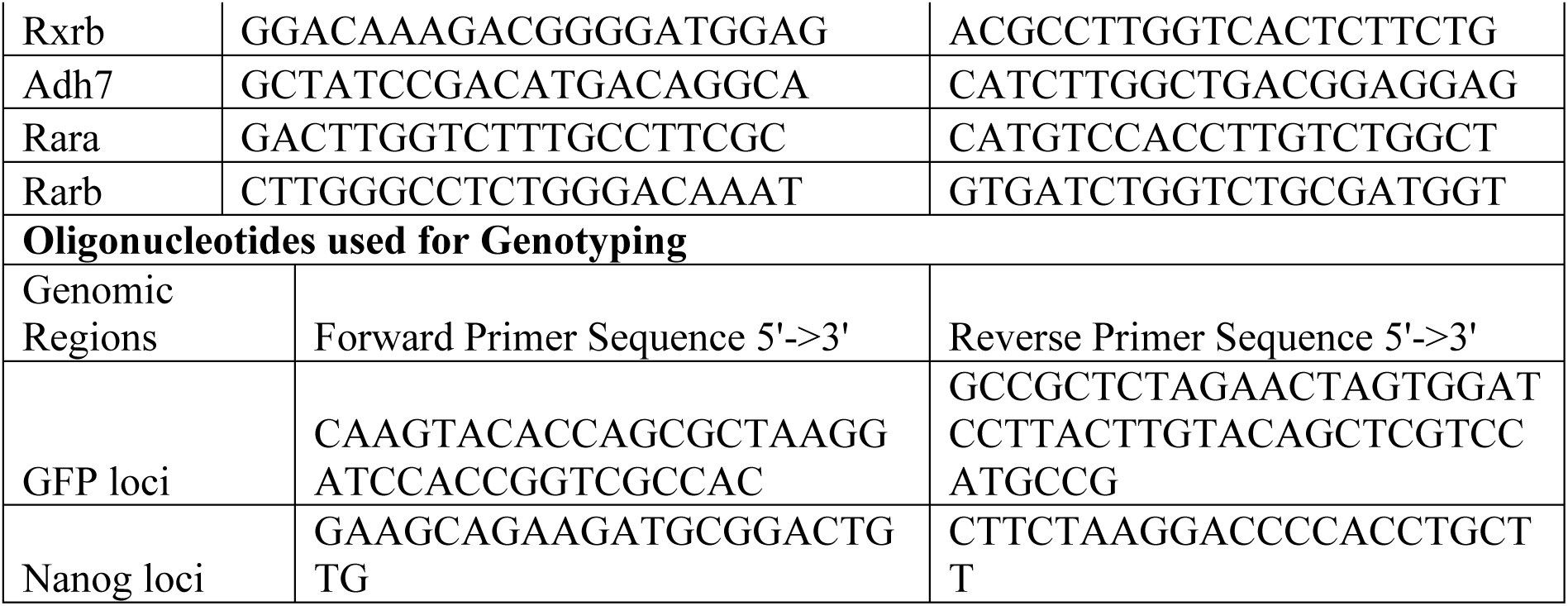

